# Transcriptomic Convergence and the Female Protective Effect in Autism

**DOI:** 10.1101/2025.01.20.634000

**Authors:** Rebecca E. Andersen, Maya Talukdar, Tyler Sakamoto, David Exposito-Alonso, Janet H.T. Song, Xuyu Qian, Seungil Lee, Nila Murugan, Ryan N. Delgado, Sijing Zhao, Gwenyth Eichfeld, Julia Harms, David C. Page, Christopher A. Walsh

## Abstract

Autism spectrum disorder (ASD) is highly heritable, and mutations in hundreds of genes have been implicated as individually rare causes of ASD^1–3^. Understanding how disruptions to these functionally diverse genes lead to the core features of ASD remains a major challenge^4^. Moreover, ASD is three- to four-fold more common in males than females^5^, and autistic females tend to carry more autosomal risk alleles for ASD compared to autistic males^6,7^, but the biological basis of this “female protective effect” (FPE) is unknown^8,9^. Here we show that individual perturbations of 18 ASD genes in human neural progenitor cells converge on shared effects on gene expression, including widespread downregulation of other ASD genes. De novo reconstruction of a gene regulatory network (GRN) enabled the identification of central transcriptional regulators, including the prominent ASD gene *CHD8* as well as novel candidates such as *REST*, that drive this transcriptomic convergence. Furthermore, the X-linked transcription factor *ZFX*, which is expressed from both the active and the inactive X chromosomes in females^10^, emerged as a key activator of many ASD genes: we propose that the higher *ZFX* expression level observed in female brain can buffer damaging mutations in diverse ASD genes, contributing to the FPE. Together, these results reveal how key GRNs can become broadly and similarly dysregulated upon disruption of individual ASD genes and provide molecular insight into the female protective effect in ASD.

## Background

Large-scale genetic studies have implicated hundreds of genes in autism spectrum disorder (ASD), with many identified through rare mutations that result in loss of function of one allele^1–3^. This suggests that gene dosage plays an important role in ASD^11^, making it critical to determine how appropriate expression levels of ASD genes are maintained. Characterization of gene regulatory networks (GRNs) could provide important insights into mechanisms mediating the proper transcriptional dosage of these ASD genes. Additionally, ASD is significantly male-biased in prevalence, and although some ASD genes have been identified on the X chromosome^12^, X-linked mutations alone cannot fully explain this sex bias^8,13^. Moreover, it has been consistently demonstrated that females require greater autosomal genetic risk to manifest ASD compared to males^6,7^, but the genetic mechanisms of this female protective effect (FPE) are unknown^8,9^.

Here, GRN reconstruction identifies key drivers of the expression of ASD genes, including *CHD8* and *REST*. These analyses also uncover the female-biased transcription factor *ZFX*, which is expressed from both the active and inactive X chromosome in females^10^, as a key regulator of ASD genes, suggesting a direct transcriptional mechanism that could contribute to the FPE. This study provides a crucial framework for uncovering how variants in diverse genes can cause convergent molecular effects that ultimately result in ASD.

## RESULTS

### Depletion of ASD Genes Reveals Widespread Transcriptomic Convergence

Functional interrogation of ASD genes revealed strongly overlapping effects on downstream targets in human neural progenitor cells (NPCs). Knockdown (KD) studies were performed for 20 ASD genes, including 18 genes from the Simons Foundation Autism Research Initiative (SFARI) Gene database^14^ and two long non-coding RNAs (lncRNAs) previously implicated in ASD (*NR2F1-AS1*^15^ and *SOX2-OT*^16^). The RNA-degrading CRISPR enzyme CasRx^17^ and an array of three guide RNAs (gRNAs) per target (**Supplementary Table 1**) were used to induce individual KD of each target gene relative to non-targeting controls (NTCs) in NPC cultures from a control human male (XY) induced pluripotent stem cell (iPSC) line (**Fig. 1a-b** and **Extended Data Fig. 1**). RNA-sequencing (RNA-seq) analysis after 24 hours of KD demonstrated that nearly all perturbations led to a substantial number of significantly differentially expressed genes (DEGs; adj. p < 0.05), with a median of 920 DEGs per perturbation (25^th^ percentile: 644, 75^th^ percentile: 1485) (**Extended Data Fig. 2** and **Supplementary Table 2**). To ensure sufficient statistical power for downstream analyses of these DEGs, we focused on the 18 perturbations (18/20, 90%) that resulted in greater than 20 DEGs, excluding the *ARHGAP5* and *ARID2* knockdown samples from further analysis. The remaining perturbations each led to a minimum of 287 DEGs, enabling robust characterization of these samples.

**Figure 1.**
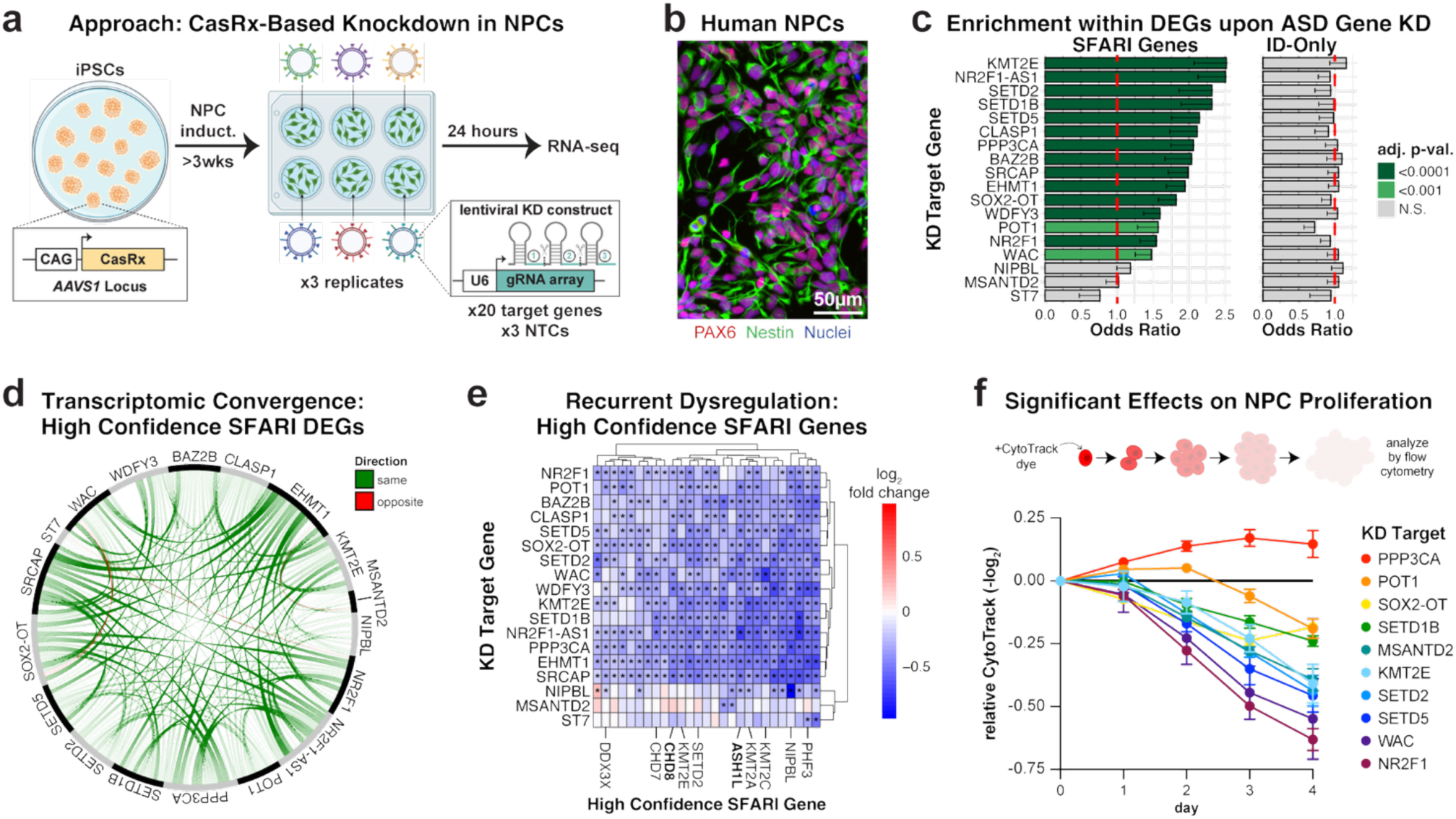
Depletion of ASD genes reveals widespread transcriptomic convergence. **a**, Experimental design schematic. iPSCs: induced pluripotent stem cells. **b,** Immunocytochemistry of iPSC-derived NPC cultures, representative of three independent analyses with similar results. **c,** Bar plot depicting enrichment of SFARI genes or ID-only genes within the DEGs of each KD condition, excluding the targeted gene itself. Bars indicate the odds ratio (OR) from a one-sided Fisher’s exact test; error bars indicate the lower bound of the 95% confidence interval on the OR (no upper bound is defined for a one-sided test). p-values were adjusted using Benjamini-Hochberg correction for multiple hypothesis testing. **d,** Circos plot depicting high confidence SFARI DEGs shared between different KD conditions (see also Extended Data Fig. 3a). **e,** Heatmap showing the relative expression of high confidence SFARI genes that are DEGs in at least 10 KD conditions (see also Extended Data Fig. 3b-c). Statistically significant DEGs were identified using DESeq2 (Wald test) with Benjamini-Hochberg correction for multiple hypothesis testing, and are indicated with *. Top ASD risk genes from ^1^ are bolded. **f,** Relative CytoTrack dye intensity (-log_2_) over time. Only KD conditions that exhibited statistically significant effects that were consistent over time are displayed. Plot depicts mean +/- standard error of the mean (SEM). Statistical significance was determined using ordinary two-way ANOVA with Dunnett’s multiple comparisons test. At least three biologically independent samples per condition were analyzed for each timepoint. See **Supplementary Table 10** for full statistical results.

Transcriptome-wide analyses revealed that ASD gene KDs caused remarkably widespread disruption of other ASD genes. Most sets of DEGs exhibited significant enrichment for known ASD genes in the SFARI gene database (15/18, 83.3%) (**Fig. 1c** and **Supplementary Table 3**). In contrast, there was no significant enrichment for genes implicated in intellectual disability (ID) but not ASD (ID-only genes) (**Fig. 1c** and **Supplementary Table 3**) (**Methods**), despite their comparable expression in NPCs (ID-only genes: 1062/1242, 85.5% expressed; SFARI genes: 797/1230, 64.8% expressed). This suggests that perturbation of ASD genes especially affects other ASD genes, as opposed to neurodevelopmental genes more broadly.

Notably, we observed a striking pattern of transcriptomic convergence across perturbations, which we define as statistically significant overlap of the same DEGs (**Extended Data Fig. 3**). This was particularly evident for high confidence SFARI (HC-SFARI) genes, the SFARI genes in the highest evidentiary tier for association with ASD (**Fig. 1d**). Moreover, HC-SFARI genes tended to shift in a concordant direction as a result of different perturbations even when they were not significant DEGs within a particular KD sample (**Extended Data Fig. 3c**). This included the prominent ASD risk gene *CHD8*^18^, which was significantly downregulated by most (11/18, 61.1%) of the KDs. Remarkably, a total of 27 HC-SFARI genes were DEGs in at least 10 of the KDs (27/183, 14.8%, p = 4.1e-9), and all were consistently downregulated (**Fig. 1e**), demonstrating convergent dysregulation of many of the strongest risk genes for ASD.

High levels of CasRx have been reported to cause non-specific RNA degradation of a magnitude that correlates with the expression level of the KD target, while lower CasRx levels avoid this issue^19^. In our NPC transcriptomic studies, CasRx was expressed only from a single safe harbor locus (*AAVS1*), limiting its expression. We also performed several additional experiments to confirm that non-specific RNA degradation was not impacting our results. Firstly, we found no relationship between the levels of expression of KD targets and the numbers of resulting DEGs either when considering the whole transcriptome (p = 0.990, R^2^ = 8e-6, linear regression) or SFARI genes specifically (p = 0.784, R^2^ = 0.0043, linear regression) (**Extended Data Fig. 4a**). We next generated NPCs expressing the fluorophores mKO2, mScarlet, and iRFP670 and targeted each individually with CasRx (**Supplementary Table 4**); flow cytometry showed no collateral effects on fluorescence intensity of non-targeted fluorophores **(Extended Data Fig. 4b** and **Supplementary Table 5**). Finally, we confirmed strong transcriptomic convergence on shared downregulated DEGs after performing additional KD studies of a subset of our target genes using short hairpin RNAs (shRNAs) in NPCs (**Extended Data Fig. 4c-d**, **Extended Data Fig. 5**, and **Supplementary Table 6**). Re-analysis of data from a study that used Cas9 to knock out ASD genes in iPSCs and iPSC-derived excitatory neurons (ExNs)^20^ similarly demonstrated strong transcriptomic convergence on shared downregulated DEGs both within each cell type and between cell types (**Extended Data Fig. 5** and **Supplementary Table 6**). Recent studies appearing since our work was submitted have also noted convergence across diverse ASD models^21,22^. Together, these data confirm that disruption of ASD genes results in transcriptomic convergence across a variety of cell types and experimental contexts.

### Transcriptomic Convergence Nominates Candidate ASD Risk Genes

The marked transcriptomic convergence we observed in our KD studies suggested that candidate ASD risk genes could be nominated by the following features: 1) consistent downregulation following ASD gene KDs, and 2) dosage sensitivity, as indicated by high genic constraint scores (**Supplementary Table 7, Methods**). This approach yielded 397 genes (**Supplementary Table 8**), including 113 SFARI genes (OR = 5.87, adj. p = 1.85e-39) and 49 HC-SFARI genes (OR = 11.17, adj. p = 1.11e-29), demonstrating that these criteria effectively prioritize known ASD genes. Notably, this set of genes showed no enrichment for ID-only genes (OR = 1.01, adj. p = 0.509). Further supporting the ASD relevance of the remaining 284 non-SFARI genes, all of them exhibit significant co-expression with SFARI genes (all adj. p < 0.05), which has previously been shown to help prioritize novel ASD risk genes^4^. Moreover, at least forty of these genes show some association with ASD in one or more major sequencing studies (**Supplementary Table 9**). For example, mutations in *AKT3*, while classically associated with brain overgrowth disorders like megalencephaly, have recently been associated with ASD; other genes such as *KIF11*, *KIF2A* and *SOS1* have also been associated with ASD in multiple studies (see references in **Supplementary Table 9**). Therefore, the remaining genes merit consideration as candidate ASD risk genes.

### ASD Genes Regulate NPC Proliferation

Many ASD gene perturbations also caused similar phenotypic effects in NPCs. Proliferation analysis using heritable dye (CytoTrack) revealed that 9/18 (50%) of the KDs led to a statistically significant decrease in proliferation (**Fig. 1f**, **Extended Data Fig. 6**, and **Supplementary Table 10**), including several genes implicated in microcephaly (*SETD1B*^23^, *SETD2*^24^, *SETD5*^25^, *KMT2E*^26^, and *WAC*^27^). In contrast, only 1/18 (5.6%) of the KDs led to a significantly increased NPC proliferation rate. This suggests that reduced NPC proliferation is a common phenotype that results from disruption of several different ASD genes, consistent with previous findings^28^. Deciphering the specific relationships between these molecular phenotypes and abnormal head size in ASD is complex, since reports implicate increased rates of both microcephaly and macrocephaly across individuals with autism^29,30^. Indeed, two KD targets that we found to decrease proliferation in NPCs – *SETD2*^24^ and *KMT2E*^26^ – have been associated with either microcephaly or macrocephaly depending on the specific variants studied in these genes. Thus, although disruption of NPC proliferation is a convergent phenotype observed across many of our perturbations, how this relates to downstream changes in head size remains to be determined.

### Disruption of ASD Genes Alters Cerebral Organoid Development

Inducible KD in cerebral organoids further revealed shared neurodevelopmental phenotypes. To enable these organoid studies, we created the FLEx-based Inducible CRISPR Knockdown (FLICK) construct. FLICK enables control over the timing of KD through doxycycline-inducible genetic recombination that results in constitutive expression of CasRx and an EGFP reporter (**Fig. 2a**), providing robust KD (**Extended Data Fig. 7a**). The FLICK constructs also contain identifying sequences that can be captured during single-cell RNA-seq (scRNA-seq) and used to determine the KD target in a given cell, enabling the analysis of mosaic cerebral organoids.

**Figure 2.**
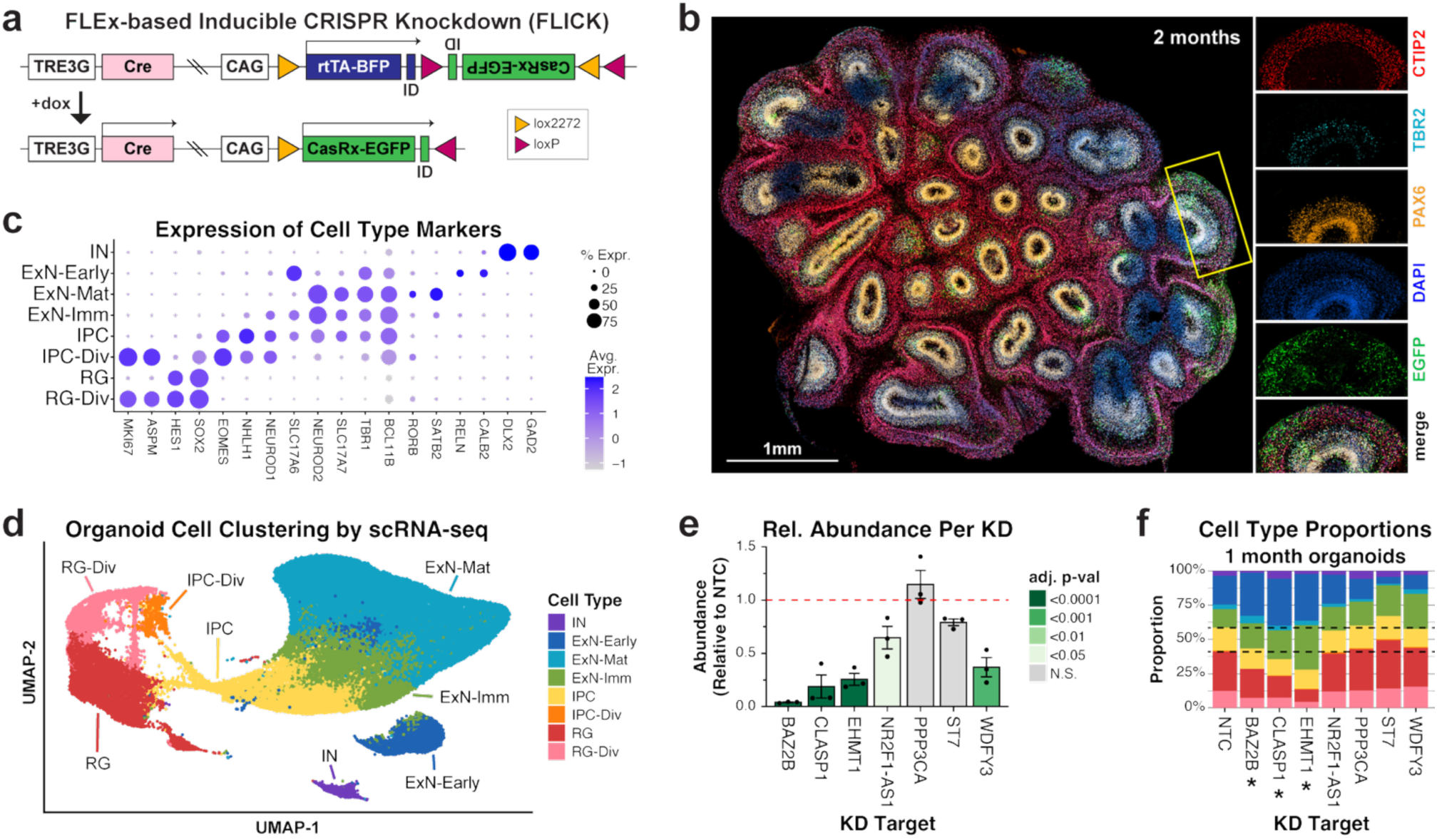
Disruption of ASD genes alters cerebral organoid development. **a**, Design of FLICK constructs for doxycycline-inducible CasRx KD. FLICK constructs also express gRNAs (not shown). **b,** FLICK organoid after 2 months of development, representative of three independent analyses with similar results. Immunohistochemistry was performed to visualize expression of the following markers: EGFP = FLICK construct; DAPI = nuclei, PAX6 = NPCs; TBR2 = intermediate progenitor cells; CTIP2 = excitatory neurons. **c,** Expression of canonical brain marker genes used for identifying cell types by scRNA-seq. **d,** UMAP of integrated long-term organoid scRNA-seq data collected at one month and two months. **e,** Relative abundance of cells for each KD target from mosaic organoids at one or two months. Results are normalized to the starting proportion for each sample at day 0 and displayed relative to NTCs from the same timepoint. Plot depicts mean +/- SEM. The FLICK constructs had been induced by treatment with doxycycline on d14-d15. Statistical significance was assessed using repeated measures one-way ANOVA with Dunnett’s multiple comparisons test. n = three biologically independent samples per condition. **f,** Cell type composition per KD target in one-month organoids. Dotted lines indicate the NTC proportions of major cell types (radial glia, intermediate progenitors, and neurons). Samples for which the ExN/RG proportion was significantly different than NTC are indicated with * (adj. p < 0.05). Statistical analysis was performed using ordinary one-way ANOVA with Dunnett’s multiple comparisons test, considering all long-term samples from one month (shown here) and two months (see Extended Data Fig. 7f).

For long-term studies, we targeted ASD genes whose KDs had not been found to impair NPC proliferation in our CytoTrack experiments (**Supplementary Table 10**). FLICK constructs targeting these genes or expressing NTC gRNAs were each separately transposed into iPSCs. We then used fluorescence-activated cell sorting (FACS) to isolate cells that had integrated the FLICK constructs based on their expression of the fluorescent reporter TagBFP. Following FACS, we pooled equal numbers of FLICK+ iPSCs for each KD target, along with the same number of NTC iPSCs, and cultured this mixed iPSC population for four days to allow them to recover before generating mosaic organoids (day 0). Recombination of the FLICK construct, which leads to constitutive expression of CasRx and an EGFP reporter, was induced in a subset of the cells (∼10%) by treatment with doxycycline from day 14-15, when the organoids predominantly consisted of PAX6+ NPCs (**Extended Data Fig. 7b**). These organoids were subsequently grown until either one or two months of development (**Fig. 2b**) and analyzed by scRNA-seq, which identified all expected cell types (**Fig. 2c-d** and **Extended Data Fig. 7c-d**) including radial glial progenitors (RGs), intermediate progenitor cells (IPCs), inhibitory neurons (INs), and ExNs. Additional scRNA-seq analysis further validated that the FLICK construct can produce efficient KD (**Extended Data Fig. 7e**).

Long-term organoid KDs revealed shared phenotypes across multiple perturbations. Several KDs, particularly those targeting *BAZ2B*, *CLASP1*, and *EHMT1*, were depleted from the organoids compared to NTCs (**Fig. 2e**), even though these KDs did not impair proliferation in NPC cultures (**Extended Data Fig. 6a** and **Supplementary Table 10**). This discrepancy between our NPC cultures and organoids suggests that long-term organoids, which support progenitor proliferation as well as differentiation, can reveal neurodevelopmental phenotypes that are not apparent in short-term NPC cultures. Interestingly, microcephaly has been associated with pathogenic variants in these genes^31–33^, consistent with the organoid phenotypes. Furthermore, these KDs also altered cell type proportions in organoids, producing relatively greater numbers of ExNs and fewer RGs (adj. p = 0.021 for *BAZ2B*, 0.018 for *CLASP1*, and 0.012 for *EHMT1*, ordinary one-way ANOVA with Dunnett’s multiple comparisons test) (**Fig. 2f** and **Extended Data Fig. 7f**). Together, this suggests that disruption of these genes may result in increased ExN differentiation at the expense of RG self-renewal. Most of the other KDs also trended toward an increased proportion of ExNs by the 2-month timepoint, suggesting this is a convergent neurodevelopmental phenotype, in line with previous observations^34,35^.

Depletion of ASD genes also resulted in prominent transcriptomic convergence in cerebral organoids (**Fig. 3a** and **Extended Data Fig. 7g**). RGs with different KDs exhibited highly overlapping DEGs, further supporting our findings from NPC cultures. Indeed, the DEGs identified in organoid RGs strongly overlapped with the DEGs identified from KD of the same target in NPC cultures (**Extended Data Fig. 7h**), suggesting that these effects on gene expression are not specific to a particular experimental context. Transcriptomic convergence was also observed within additional cell types including IPCs as well as ExNs (**Fig. 3a** and **Extended Data Fig. 7g**), demonstrating that the convergence of ASD genes on shared downstream effects is not unique to neural progenitors. This is consistent with studies identifying shared ASD DEGs in postmortem brain tissue from individuals with distinct causes of ASD^36–38^ as well as with widespread observations of convergent ExN phenotypes such as disordered synaptic structure or activity upon disruption of distinct ASD genes^20^, suggesting that convergence does not reflect merely a shared alteration of cell type or fate of progenitor cells. Furthermore, we find that detection of overlapping DEGs using scRNA-seq is heavily dependent upon the number of cells sequenced for each perturbation, as demonstrated by down-sampling analysis (**Fig. 3b**). This sample-size dependence may partially explain the lack of robust transcriptomic convergence observed at the individual gene level in some previous studies with smaller sample sizes^34,39^. Our use of an isogenic background also likely contributed to our detection of transcriptomic convergence, as different genetic backgrounds have been found to significantly affect neurodevelopmental phenotypes resulting from the disruption of ASD genes in organoids^34^. Finally, while some convergently disrupted genes were cell type-specific, many genes were consistently disrupted across multiple cell types, including SFARI DEGs (**Fig. 3c**). This suggests that despite cell-type-specific features impacting the transcriptomic consequences of ASD gene perturbations, certain effects could be broadly shared by diverse cell types.

**Figure 3.**
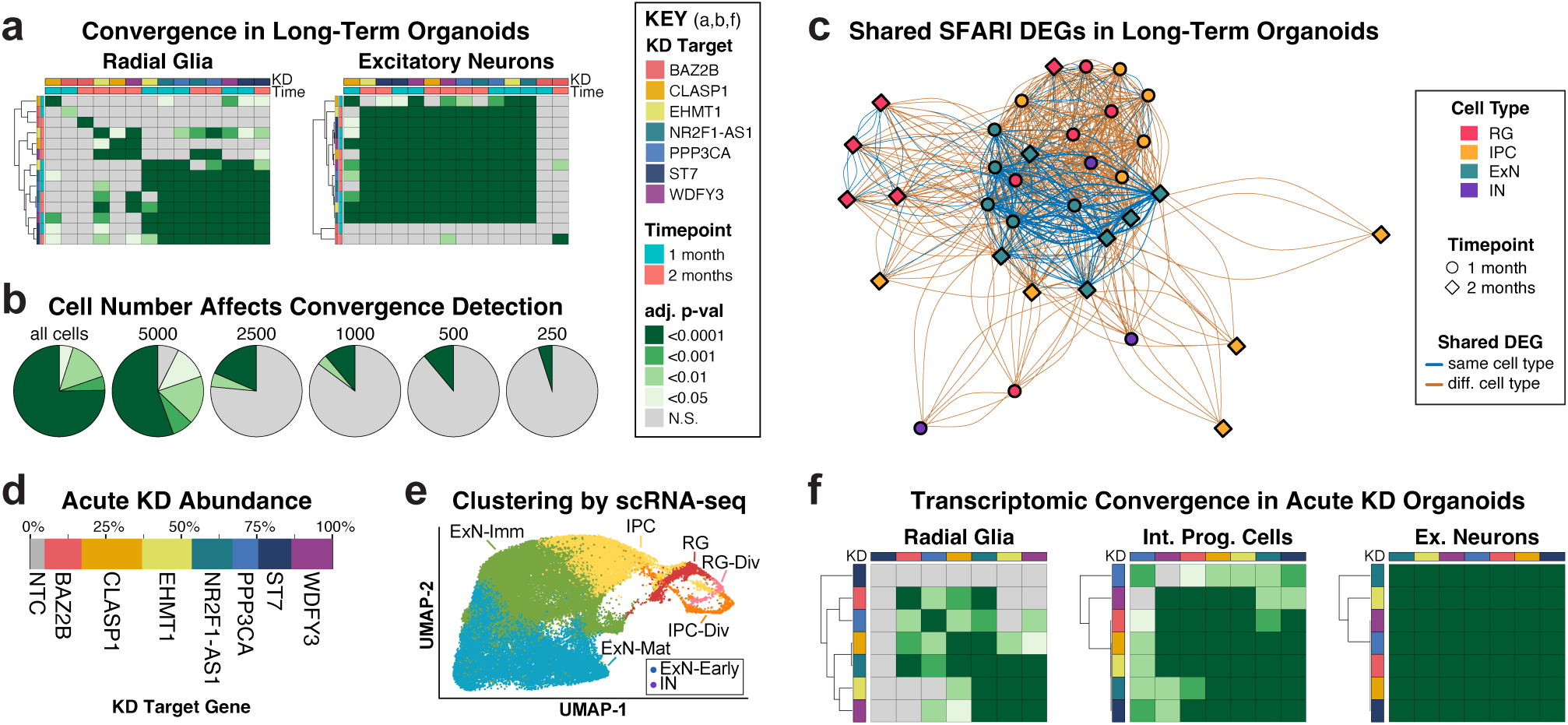
Transcriptomic convergence upon ASD gene perturbations in cerebral organoids. **a**, Heatmap of cross-enrichment of DEGs for radial glia and excitatory neurons in long-term organoids. **b,** Pie charts denoting proportions of pairwise sets of DEGs that show significant cross-enrichment (transcriptomic convergence). Down-sampling the number of cells per condition to the number indicated above each pie chart decreases the transcriptomic convergence observed. **c,** Network visualization of DEGs shared amongst different KDs conditions and cell types. The thickness of edges represents the number of shared DEGs. Nodes plotted based on clustering with the Fruchterman-Reingold algorithm (*igraph*). Nodes that are closer together have more shared DEGs on average. **d,** Proportion of cells with each KD target from scRNA-seq of mosaic organoids with acute KD. **e,** UMAP of acute KD mosaic organoids from scRNA-seq. **f,** Heatmap of cross-enrichment between different KD samples in acute KD organoids.

### Transcriptomic Convergence Following Acute Perturbations in Cerebral Organoids

Acute disruption of ASD genes in 2-month organoids revealed even greater convergence on shared DEGs. An additional set of mosaic FLICK organoids was grown for two months prior to inducing KD with doxycycline treatment from day 61-62 followed by scRNA-seq two days later, allowing us to determine the acute effects of disrupting these genes within organoids that had developed normally. This approach resulted in similar numbers of cells per KD (**Fig. 3d** and **Extended Data Fig. 7i**), enabling deeper analysis of the KDs that had been depleted from the organoids with long-term KD (**Fig. 2e**). As expected, these organoids contained a mixture of cell types (**Fig. 3e** and **Extended Data Fig. 7j**), with ExNs by far the most abundant cell type at this developmental timepoint (**Extended Data Fig. 7k**). Interestingly, many KDs resulted in ExN DEGs that were enriched for the DEGs identified from NPC cultures (**Extended Data Fig. 7l**), again suggesting shared changes in gene expression across cell types upon KD of ASD genes. Furthermore, acute KD of the target genes that had been depleted from organoids over the course of long-term KD (**Fig. 2e**) allowed us to obtain much greater cell numbers for these perturbations, revealing that these KDs also result in strong transcriptomic convergence (**Fig. 3f**).

### Key Transcription Factors, Including *REST*, Target Recurrently Dysregulated Genes

Transcriptional regulators that could mediate transcriptomic convergence in ASD were identified through MAGIC analysis (**Methods**), which uses chromatin immunoprecipitation sequencing (ChIP-seq) data from ENCODE to identify transcription factors and cofactors whose binding targets are enriched within a gene set of interest. This uncovered transcriptional regulators whose target genes were recurrently differentially expressed across many of the ASD gene KDs in NPCs (**Extended Data Fig. 8a** and **Supplementary Table 11**), including HC-SFARI genes such as *CHD2* and *SIN3A*. Of the recurrent MAGIC regulators not previously implicated in ASD, several were both highly constrained (**Supplementary Table 7)** and co-expressed with known ASD genes (**Extended Data Fig. 8a**), features demonstrated to be predictive of novel ASD risk genes^4^.

MAGIC analysis also revealed consistent drivers of convergence in organoids (**Extended Data Fig. 8a** and **Supplementary Table 11**). Of the top recurrent transcriptional regulators identified in NPCs, half (13/26, 50%) were also identified in organoid RGs (**Extended Data Fig. 8b**). MAGIC also identified recurrent ExN regulators shared across developmental stages and across long-term vs acute KD timescales, including many SFARI genes. Interestingly, nine of these genes were also identified as top transcriptional regulators in neural progenitors, suggesting that they are key drivers of convergence in multiple cellular contexts.

Of particular interest, the RE1 silencing transcription factor (REST) was significant by MAGIC in 17/18 (94.4%) NPC perturbations (**Extended Data Fig. 8a**) and consistently emerged as a top MAGIC regulator across all tested conditions (**Extended Data Fig. 8b**). While direct mutations in *REST* have not previously been associated with ASD, *REST* has critical neurodevelopmental roles and functions as a master negative regulator of neurogenesis^40,41^. Dysregulation of *REST* has also been reported following the disruption of ASD genes, such as *CHD8*^42^ and the Rett syndrome gene *MECP2*^43,44^. Re-analysis of human and mouse REST ChIP-seq binding sites (**Methods**) identified strong enrichment of SFARI genes (adj. p = 3.6e-67 and 7.7e-50, respectively), including HC-SFARI genes (adj. p = 2.2e-13 and 3.7e-8, respectively), within REST targets (**Extended Data Fig. 8c**), supporting a key role for REST in regulating ASD-relevant gene expression.

### Reconstruction of a Gene Regulatory Network Reveals Highly Interconnected ASD Genes

De novo GRN reconstruction from our NPC RNA-seq data using ARACNe enabled associating additional transcription factors and chromatin modifiers – together referred to as “regulators” – with their downstream targets (regulons) (**Methods**). We validated this approach by separately constructing “held-out” GRNs that excluded data from samples in which the KD target itself was a potential regulator, thereby preventing circularity in regulon inference. Regulons predicted from these held-out GRNs strongly overlapped the DEGs observed following KD of the corresponding target genes in NPCs (adj. p for each < 3.98e-116, combined p = 4.08e-937) (**Fig. 4a** and **Extended Data Fig. 9a**). Thus, ARACNe effectively captures bona fide regulatory relationships.

**Figure 4.**
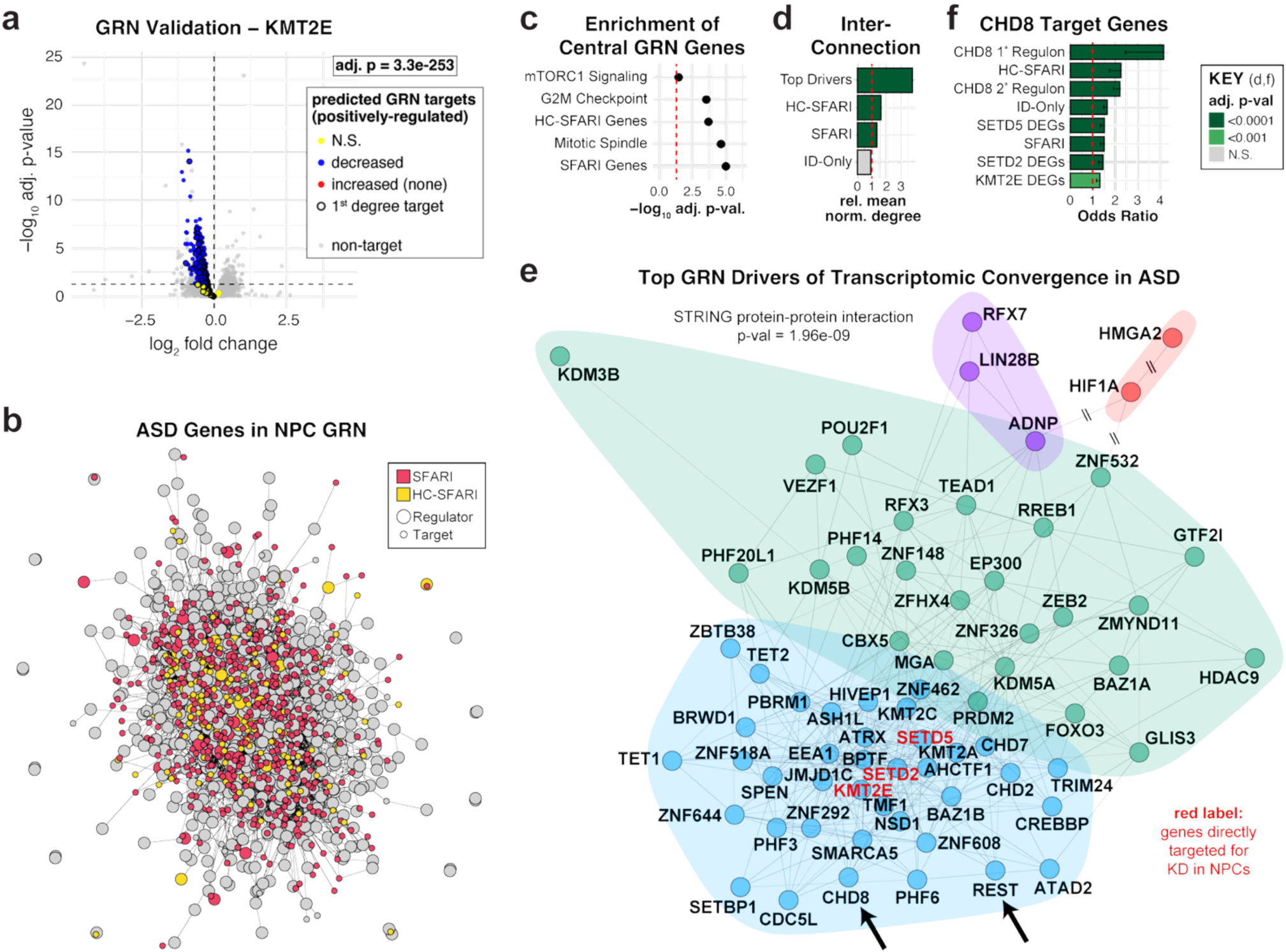
Gene regulatory network analysis identifies *CHD8* as a top driver of convergence in ASD. **a**, Volcano plot depicting differential expression upon KD of KMT2E. Genes predicted by GRN analysis to be positively regulated by KMT2E are highlighted, with statistically significantly decreased DEGs shown in blue, increased DEGs shown in red (none), and targets with no significant change shown in yellow. Statistically significant DEGs were identified using DESeq2 (Wald test) with Benjamini-Hochberg correction for multiple hypothesis testing. Statistically significant enrichment of 2nd degree regulon genes within the downregulated DEGs is indicated by the adjusted p-value (one-sided Fisher’s exact tests with Benjamini-Hochberg correction for multiple hypothesis testing). **b,** Depiction of a subset of the NPC GRN constructed using ARACNE. All regulators are shown along with first degree target genes that are SFARI genes. **c,** Gene set enrichment analysis (GSEA) (Weighted Kolmogorov-Smirnov running-sum statistic, permutation test, with Benjamini-Hochberg correction for multiple hypothesis testing) demonstrating enrichment of genes with high PageRank scores within the gene sets shown. **d,** Bar graph depicting mean normalized degree, a measure of interconnectedness, across regulators from different gene sets. **e,** Depiction of modules of top regulators identified through iterative walktrap clustering. Hashed lines indicate that those lines are not to scale. Genes of particular interest are highlighted with large arrows. KD targets that are further analyzed in panel E are highlighted in red font. **f,** Bar plot depicting enrichment of CHD8 targets within several gene sets. Data from CHD8 ChIP-seq in NPCs from Sugathan et al. (2014)^45^. Bars indicate the odds ratio (OR) from a one-sided Fisher’s exact test; error bars indicate the lower bound of the 95% confidence interval on the OR (no upper bound is defined for a one-sided test). p-values were adjusted using Benjamini-Hochberg correction for multiple hypothesis testing.

The full NPC GRN revealed a high degree of interconnection between ASD genes and their upstream regulators (**Fig. 4b**). Indeed, ASD genes were strongly enriched within central genes of the NPC GRN (adj. p = 9.81e-6 for SFARI genes, 1.99e-4 for HC-SFARI genes, centrality assessed by *PageRank* algorithm, **Methods**) (**Fig. 4c**). In contrast, ID-only genes did not exhibit enrichment within the central genes of the GRN (adj. p = 0.19), consistent with the lack of enrichment that we had observed within the DEGs resulting from ASD gene KDs (**Fig. 1c**). Additionally, central GRN genes were enriched for pathways associated with the G2M checkpoint (adj. p = 6.0e-4) and mitotic spindle (adj. p = 3.6e-5), consistent with the NPC proliferation phenotypes we had observed (**Fig. 1f** and **Supplementary Table 10**).

### GRN Analysis Identifies *CHD8* as a Top Driver of Convergence in ASD

We next identified GRN regulators that act as key drivers of ASD-associated transcriptomic convergence. This analysis revealed 70 top drivers of convergence whose activity was consistently altered by ASD gene KD (**Methods**), and whose targets are both enriched for SFARI genes and central to the GRN (**Supplementary Table 12**). Nearly half of these top drivers of convergence are SFARI genes (31/70, OR = 6.35, adj. p = 3.9e-11), and nearly one third are HC-SFARI genes (22/70, OR = 10.61, adj. p = 6.7e-12). In contrast, only seven of these top drivers of convergence are ID-only genes (7/70, OR = 2.06, adj. p = 0.078). Moreover, these top drivers of convergence were consistently downregulated across nearly all ASD gene KDs (**Extended Data Fig. 9b**). They are also highly interconnected in the GRN (p < 1e-6) (**Fig. 4d**), as well as the STRING database of protein-protein interactions (p = 2.0e-9) (**Methods**). Taken together, this suggests that these genes not only act as central regulators of many ASD genes but also cross-regulate each other, driving transcriptomic convergence in ASD.

Iterative walktrap clustering identified four gene modules within the 70 top drivers of convergence (**Fig. 4e**), one of which (blue) was especially strongly interconnected. This module exhibited very high cross-interaction through STRING analysis (p < 1e-16) and included *REST* (**Extended Data Fig. 8**) and several notable ASD genes including *CHD8*, one of the strongest known risk genes for ASD^18^. Importantly, the inferred *CHD8* regulon from the GRN was strongly enriched (1^st^ degree regulon adj. p = 4.6e-7, 2^nd^ degree regulon adj. p = 3.9e-20) for direct CHD8 target genes previously identified through ChIP-seq from NPCs^45^ (**Fig. 4f**), further supporting the validity of ARACNe-predicted regulatory relationships. Moreover, consistent with previous reports^45,46^, CHD8 ChIP-seq targets were also highly enriched for NPC-expressed SFARI genes (adj. p = 2.6e-8) and HC-SFARI genes (adj. p = 8.5e-8). The *CHD8*-containing module also included three transcriptional regulators (*SETD2*, *SETD5*, and *KMT2E*) that we had directly perturbed, and DEGs from these KDs exhibited significant enrichment for CHD8 ChIP-seq targets (adj. p = 1.2e-5, 7.8e-8, and 2.6e-4, respectively) (**Fig. 4f**), further demonstrating convergent regulation of shared targets by critical ASD genes. This analysis highlights *CHD8*, a high confidence ASD risk gene, as a key driver of transcriptomic convergence in ASD.

### The X-Linked Transcription Factor ZFX is a Female-Biased Regulator of ASD Genes

Prior studies have found that genes downregulated in ASD tend to be more highly expressed in the neurotypical female versus male brain^50,51^, raising the possibility that this baseline female-biased expression may contribute to the FPE^8,50,51^. We identified 82 genes that were consistently downregulated by ASD gene KD and exhibited female-biased expression in the brain (**Methods**), 28 of which were highly constrained (**Extended Data Fig. 10a**), including 12 highly constrained SFARI genes (OR = 2.79, p = 8.65e-3) (**Fig. 5a**). Re-analysis of single-nucleus RNA-sequencing data (**Methods**) from the frontal cortex of neurotypical male controls and autistic males – including individuals with 15q duplication syndrome and genetically idiopathic ASD – revealed that many of these highly constrained genes are significantly downregulated in excitatory neurons from autistic males, further supporting their relevance to ASD (**Fig. 5a** and **Extended Data Fig. 10a**).

**Figure 5.**
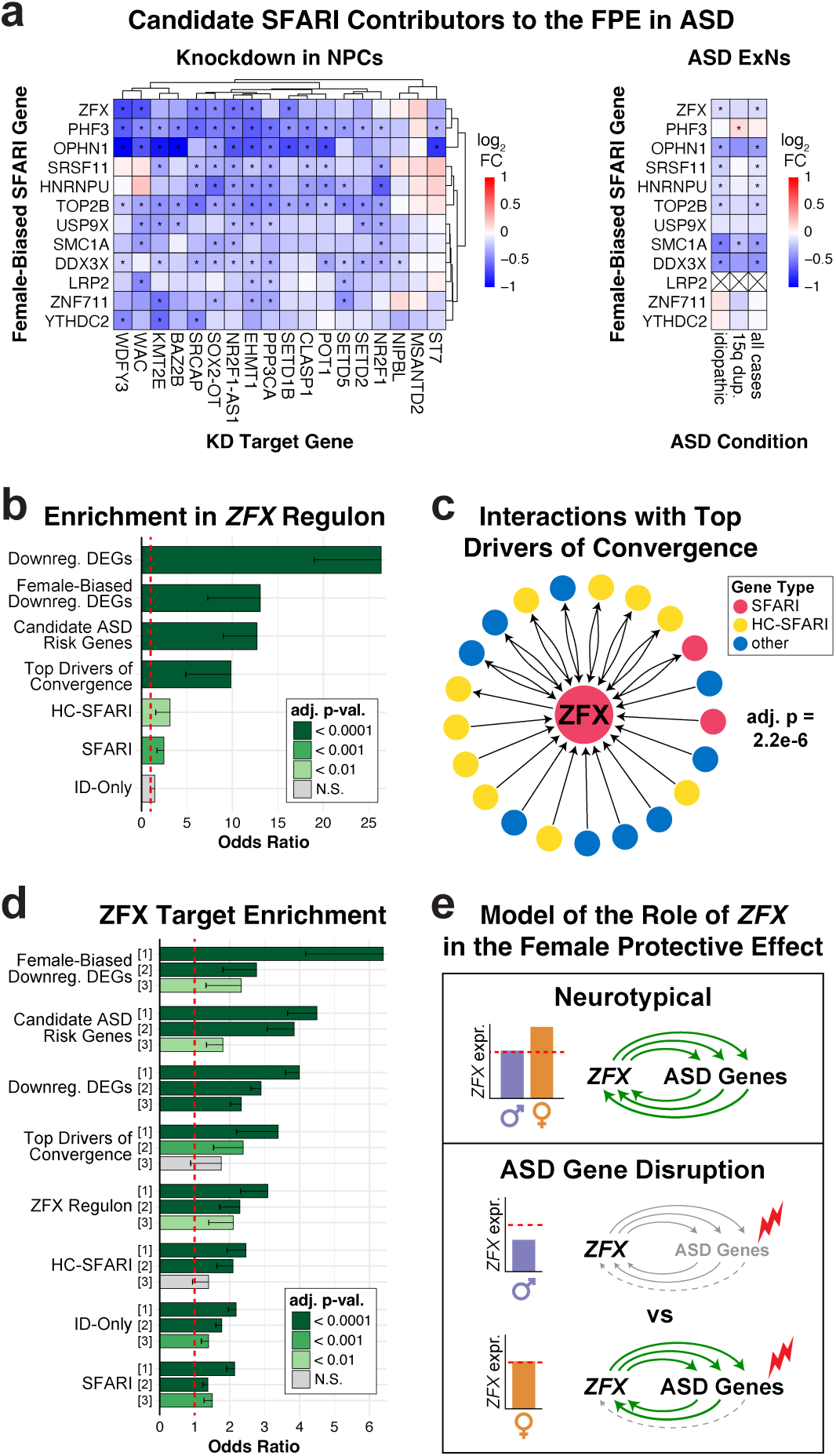
The X-linked transcription factor ZFX is a female-biased regulator of ASD genes. **a**, Heatmaps showing the relative expression of SFARI genes that are candidate contributors to the FPE in ASD. Left: expression in NPCs across KD conditions relative to NTCs, with statistically significant DEGs (indicated with *) identified using DESeq2 (Wald test) with Benjamini-Hochberg correction for multiple hypothesis testing. Right: expression in ExNs from males with ASD relative to male idiopathic controls, re-analyzed from Dias et al. (2024)^47^, with statistically significant DEGs (indicated with *) identified using MAST (likelihood ratio test) with Bonferroni correction for multiple hypothesis testing. Lack of detection is indicated with X. See also Extended Data Fig. 10a. **b,** Bar plot depicting enrichment of ASD-relevant gene sets within the *ZFX* regulon, as identified by GRN analysis. Bars indicate the odds ratio (OR) from a one-sided Fisher’s exact test; error bars indicate the lower bound of the 95% confidence interval on the OR (no upper bound is defined for a one-sided test). p-values were adjusted using Benjamini-Hochberg correction for multiple hypothesis testing. **c,** Depiction of regulatory relationships between *ZFX* and the top drivers of convergence in ASD (previously identified in Fig. 4e), as identified by GRN analysis. Statistically significant enrichment of the top drivers of convergence within the *ZFX* regulon is indicated by the adjusted p-value (one-sided Fisher’s exact tests with Benjamini-Hochberg correction for multiple hypothesis testing). **d,** Bar plot depicting enrichment of ASD-relevant gene sets within the following *ZFX* target sets: [1] targets in C4-2B cells, identified in San Roman et al. (2024)^48^; [2] common targets identified in Rhie et al. (2018)^49^; and [3] targets in MCF-7 cells, identified in San Roman et al. (2024)^48^. Bars indicate the odds ratio (OR) from a one-sided Fisher’s exact test; error bars indicate the lower bound of the 95% confidence interval on the OR (no upper bound is defined for a one-sided test). p-values were adjusted using Benjamini-Hochberg correction for multiple hypothesis testing. **e,** Schematic depicting a model of the contribution of *ZFX* to the FPE in ASD.

Among these, *ZFX* is of particular interest because it is expressed from both the active and inactive X chromosome in females^10^, making it one of the most highly and consistently female-biased genes across diverse tissues^52^, including the developing cerebral cortex (**Extended Data Fig. 10b-c**). Moreover, a recent report implicated pathogenic *ZFX* variants in ASD^53^, and additional coding variants in the *ZFX* gene have also been identified in ASD probands^3,54,55^. *ZFX* encodes a potent transcriptional activator of genome-wide targets^48^, and we identified many regulatory relationships between *ZFX* and ASD-relevant gene sets in our GRN (**Fig. 5b**), including the top drivers of transcriptomic convergence in ASD we had previously identified (OR = 9.8, adj. p = 2.2e-6). We also found that *ZFX* is a downstream target of many of these top drivers, with several reciprocal regulatory relationships (**Fig. 5c**). The strong enrichment of ZFX targets for ASD-relevant genes was further confirmed by re-analysis of ZFX ChIP-seq data (**Fig. 5d**). As *ZFX* is recurrently downregulated upon disruption of ASD genes, its female-biased expression in combination with its role as a regulator of ASD-relevant genes could result in cascading and broad sex-differential impacts on ASD gene expression, potentially contributing to the FPE in ASD (**Fig. 5e**).

## DISCUSSION

Here, we show widespread transcriptomic convergence following depletion of ASD genes and highlight key chromatin modifiers and transcription factors, including *CHD8* and *REST*, as critical drivers of this convergence. These drivers were highly interconnected with each other, often via mutual regulatory relationships, revealing how disruption of a single regulator can lead to changes that propagate throughout a GRN. We also identified the transcription factor *ZFX*, which is expressed from both the active and inactive X chromosome in females, as a key transcriptional regulator of ASD genes; the female-biased expression of *ZFX* could thus buffer the effects of ASD gene disruptions in females, contributing to the FPE in ASD. Taken together, this work provides a model for how diverse genes can converge on core transcriptomic changes and ultimately generate shared ASD phenotypes, as well as how sex-differential activity of these genes can lead to male-biased risk for ASD.

Our ability to detect widespread transcriptomic convergence likely benefited from several experimental design choices that were intended to minimize variability. For instance, to avoid confounding effects of different genomic backgrounds, we opted to perturb the ASD genes in an isogenic background. Additionally, we performed these perturbations by introducing KD constructs in a polyclonal manner instead of creating different cell lines for each ASD target gene, as the process of clonal selection can introduce line-to-line variability^56^. For our cerebral organoid studies, we created the FLICK KD construct so that we could generate mosaic organoids containing all KD conditions as well as the NTCs, minimizing any effects from organoid-to-organoid variability. By reducing variability across our experimental designs, we likely enhanced our sensitivity to detect convergence.

We also optimized our RNA-seq approaches for the different experimental contexts. Initially, we used highly homogeneous NPC cultures, enabling us to perform bulk RNA-sequencing on this single cell type. This allowed deep transcriptomic profiling, avoiding the issues of “dropout” that affect scRNA-seq^57^. However, for samples that contain a heterogeneous mix of different cell types, such as cerebral organoids, bulk RNA-seq could reflect not only changes in gene expression but also differences in relative cell type abundances. Thus, we used scRNA-seq to analyze our organoids, and we sought to overcome the decreased sensitivity of this approach by sequencing a large number of cells for each KD condition. As our down-sampling analysis demonstrated (**Fig. 3b**), this large cell number was critical for detecting transcriptomic convergence through scRNA-seq. Furthermore, for our NPC experiments we performed RNA-seq 24 hours after introducing the KD constructs. We selected this short timeframe in order to focus our analysis on the initial effects of each KD, rather than long-term consequences that could be more heavily impacted by compensatory responses and stochastic effects. This short timeframe may be particularly well suited for detecting transcriptomic convergence, which further benefited our GRN reconstruction.

Building an understanding of the complex relationships within GRNs is critical for identifying key “rate-limiting” genes that affect the GRN as a whole and ultimately lead to a particular biological outcome. Prior studies of Hirschsprung disease have identified an analogous GRN centered on the rate-limiting gene *RET*, which can be impacted by direct mutations in *RET* itself or disruptions of upstream regulators of *RET*^58^. Here, we have similarly identified critical genes, such as *CHD8* and *REST*, whose effects can propagate throughout the GRN and drive transcriptomic convergence in ASD. Our findings are consistent with prior studies identifying CHD8 as an important regulator of ASD gene expression^45,46,59^. Moreover, haploinsufficiency of *CHD8* has been found to result in dysregulation of *REST* in the mouse brain^42^, consistent with the highly interconnected relationships we found within our GRN. Additionally, disruption of the Rett syndrome gene *MECP2* affects REST^43,44^, and *REST* is dysregulated in ASD brain tissue relative to neurotypical controls^42^, further supporting its relevance to ASD. Given that REST is a master negative regulator of neuronal genes^40,41^ and that CHD8 affects proliferation and neurogenesis^60^, it is plausible that the convergent dysregulation of CHD8/REST targets could directly underlie the altered proliferation we found in NPCs (**Fig. 1f**) as well as the increased relative abundance of excitatory neurons that we observed following some of the ASD gene KDs in organoids (**Fig. 2f**). Several prior studies also identified imbalances in neural proliferation versus differentiation following ASD-relevant perturbations^28,60,61^; thus, future work to further characterize the molecular mechanisms linking our observed convergent changes in gene expression to these shared neurodevelopmental phenotypes will be highly valuable.

Our results also provide new insights into the male bias of ASD. Downregulation of *ZFX* by KD of many different ASD genes in NPCs as well as lower *ZFX* levels in excitatory neurons from individuals with ASD (**Fig. 5a**) suggest that decreased *ZFX* expression may be a recurrent transcriptional feature of ASD. *ZFX*’s higher baseline level in the female brain due to its expression from both X chromosomes could thus buffer females against this ASD-associated downregulation. Consistent with this, a recent study of sex-biased gene expression across cell types in the cortex found that promoters of autosomal genes exhibiting female-biased expression were enriched for ZFX transcription factor motifs, while promoters of male-biased genes were not^62^. While males have a Y-linked homolog of *ZFX* called *ZFY*, it encodes a weaker transcriptional activator than *ZFX*^48^, and *ZFY*’s GRN regulon showed no enrichment for SFARI genes (OR = 1.1, adj. p = 0.36), suggesting divergent roles with respect to ASD. Further functional characterization of *ZFX* and other genes expressed from the inactive X chromosome may provide critical insights into how female-biased expression may impact ASD risk^13^. Moreover, it will also be valuable to directly compare the transcriptional effects of ASD gene perturbations between male cell lines, as used in this study, and female cell lines. Indeed, our use of a male rather than female founder cell line may have contributed to the striking changes in gene expression we observed, as female cells may be able to better tolerate the same ASD gene KDs due to the higher baseline expression of *ZFX* and/or additional factors that contribute to the FPE.

Given that many genes implicated in ASD show rare heterozygous loss-of-function (LoF) mutations^1^, our study prioritized ASD genes with a high probability of loss of function intolerance. Additionally, we focused on genes that exhibit a “head-to-head” configuration with an adjacent lncRNA; several lncRNAs in such a configuration have been found to fine-tune the expression of their neighbors^63^, potentially identifying particularly dosage-sensitive genes. We additionally included two lncRNAs that have themselves been implicated in ASD through heterozygous LoF mutations^15,16^. Our Cas13-based knockdowns on average produced a 48.2% decrease in expression of the target genes (**Supplementary Table 2**), an experimental approach that appears well suited for studying how heterozygous LoF of these haploinsufficient genes can contribute to ASD.

While our perturbations focused on genes associated with ASD through rare single-gene mutations, it is likely that other classes of ASD-associated variants also disrupt this same convergent network. Supporting this, studies of postmortem brain tissue have demonstrated that idiopathic ASD cases share core transcriptomic changes^38^ and overlapping DNA methylation signatures^64^ with cases of ASD caused by rare mutations of large effect size, such as chromosome 15q11-13 duplication. Moreover, reports suggest that common alleles of smaller effect size implicated in ASD can also drive changes that converge with those caused by rare ASD variants^65^. This raises the intriguing possibility that distinct causes of ASD – including rare single-gene mutations, large copy number variants (CNVs), and common genetic variation – could result in similar phenotypes by each converging on disruption of shared ASD-relevant GRNs. Further characterization of these GRNs has broad potential to clarify our understanding of the genetic architecture of ASD across a wide variety of genetic variant classes, uncovering potential therapeutic targets that could benefit individuals with diverse causes of ASD. More broadly, while this work focused on ASD, the approaches used here are widely applicable to different cell types and conditions. Thus, this study serves as a framework for uncovering critical transcriptional regulators across a wide diversity of contexts, from development through disease.

## MATERIALS AND METHODS

### Knockdown with CasRx

CasRx^17^, specifically NLS-RfxCas13d-NLS (Addgene #109049), was used in combination with target-specific guide RNAs (gRNAs) to knock down (KD) targeted transcripts. A deep-learning-based tool^70^ was used to design gRNA spacer sequences targeting the genes of interest. These spacer sequences were filtered to exclude homopolymers (greater than 4 As, greater than 3 Cs, Gs, or Ts). Spacer sequences with potential off-target interactions were identified using BLAST^71^ and excluded. For each target gene, 3 separate spacer sequences were selected and combined into a tandem pre-gRNA array, with each spacer sequence preceded by the CasRx direct repeat (DR36) sequence^70^: CAAGTAAACCCCTACCAACTGGTCGGGGTTTGAAAC. Tandem pre-gRNA arrays are processed by CasRx into individual gRNAs for KD^17^. See **Supplementary Table 1** for full sequences.

### Molecular Cloning

To create constructs for the initial NPC KD experiments, the lentiviral pSicoR plasmid^72^ (Addgene #11579) was used as the backbone. The mouse U6 promoter was replaced with the human U6 promoter, which was used to drive expression of the CasRx gRNA arrays. These constructs were used in combination with a CasRx knock-in cell line (described below).

To obtain enough NPCs for CasRx KD validation by auto-western blot (described below), an all-in-one construct was created that contained CasRx and EGFP expressed from the CAG promoter element and gRNA arrays expressed from the human U6 promoter. The backbone also contained inverted terminal repeats (ITRs), allowing the construct to be integrated into the genome by piggyBac transposase^73^. These constructs were based on the LiOn design^74^ (Addgene #154019), such that expression of CasRx and EGFP was dependent upon recombination of the construct by piggyBac transposase, preventing expression from the episomal plasmid. This all-in-one construct was also used for the fluorophore KD assays and NPC proliferation assays.

To express fluorophores in NPCs, iOn-based constructs were created with mKO2, mScarlet, or iRFP670 driven by the CAG promoter element.

For shRNA-based KD, design of shRNAs was performed using the Horizon siDesign Center tool (https://horizondiscovery.com/en/ordering-and-calculation-tools/sidesign-center). Predicted shRNAs starting with G were selected, and those with potential off-target interactions were identified using BLAST^71^ and excluded. The selected shRNAs (**Supplementary Table 13**) were cloned into the pSicoR plasmid^72^ (Addgene #11579).

For organoid experiments, we created the FLEx-based Inducible CRISPR Knockdown (FLICK) construct by engineering several features into a backbone incorporating the FLEx design^75^. In particular, we included a modified Cre recombinase that includes an intron, which prevents recombination of the plasmid during cloning^76^. We also included CasRx^17^ as well as all necessary components of the Tet-On 3G system^77^ to enable doxycycline-inducible recombination. Furthermore, we included identifying barcode sequences shortly upstream of the polyA sites for the fluorescent reporters, which enabled determination of which KD construct was incorporated in each cell through single-cell RNA sequencing (scRNA-seq). Finally, we incorporated ITRs, allowing the construct to be integrated into the genome by piggyBac transposase^73^. The sequences of all constructs were verified by whole-plasmid sequencing (Plasmidsaurus).

### Cell Culture

All human cell culture work was performed under protocols approved by Boston Children’s Hospital IRB #S07-02-0087. The control male human iPSC line “280” was previously generated and fully characterized^78^. Human iPSCs and iPSC-derived NPCs were grown on tissue culture plates pre-coated with Matrigel (Corning #354277) or Geltrex (ThermoFisher #A1413302). The cells were passaged as needed using Accutase (Millipore Sigma #SCR005) or ReleSR (StemCell Technologies #100-0483). Human iPSCs were cultured in mTeSR™ Plus medium (StemCell Technologies #100-0276). Human NPCs^79^ were generated from iPSCs using the STEMdiff™ SMADi Neural Induction Kit (StemCell Technologies #08582) by following the Monolayer Culture Protocol for at least three weeks. NPCs were continuously grown in Neural Induction Medium + SMADi and were not switched to Neural Progenitor Medium. The medium was supplemented with 10uM Y-27632 (StemCell Technologies #72304) for approximately 1 day following passaging of iPSCs or NPCs.

Cerebral organoids were generated from 280-iPSCs using the STEMdiff™ Cerebral Organoid Kit (StemCell Technologies # 08570) with some recommendations from Giandomenico et al. (2021)^80^ as well as additional modifications. In particular, the organoids were embedded in Matrigel (Corning #354277) at day 7 and grown in Expansion Medium until day 14, at which point they were removed from Matrigel and switched to Maturation Medium on an orbital shaker. Please see full details on protocols.io: https://dx.doi.org/10.17504/protocols.io.x54v95p5ql3e/v1. None of the cell lines were authenticated. All cell lines routinely tested negative for mycoplasma contamination.

### Derivation of CasRx Knock-in iPSCs

CasRx^17^ was knocked into the *AAVS1* safe harbor locus of 280-iPSCs using targeting constructs that were previously described^77^. Individual iPSC colonies were picked and expanded. Southern blot analysis was performed by Celplor to confirm correct targeting of the *AAVS1* locus. Karyotype analysis was performed using KaryoStat (ThermoFisher), which revealed a 2.4Mb gain on chromosome 1, which is described in microarray nomenclature as arr[GRCh37] 1q32.1(204108163_206539638)x3. This region did not contain any of the target genes.

### Lentivirus

Lentivirus was generated using standard protocols that have previously been described^81^. Lentiviruses were applied at a multiplicity of infection (MOI) of 4.0, as this MOI yields >98% transduced cells. RNA was extracted 24 hours later.

### Library Preparation for RNA Sequencing (RNA-Seq)

Bulk RNA was extracted using the Direct-zol RNA Microprep Kit (Zymo Research #R2062), and 3’ RNA-seq libraries were prepared following a modified version of the TagSeq protocol^82^, incorporating recommendations and pooling strategies from the CheapSeq protocol^83^. Briefly, 200ng of total RNA from each sample was combined with an indexed oligo(dT) primer and buffer (from NEB #M0466L) as described for CheapSeq. RNA was fragmented in a pre-heated thermocycler at 94C for 2.5 min without the lid, after which the samples were moved immediately to ice. Each sample was then mixed with a template-switching oligo (TSO) that contained an index and a unique molecular identifier (UMI) along with additional reagents required for reverse transcription, as described for CheapSeq. The reverse transcription reaction was performed at 42C for 90 minutes, without heat inactivation. Remaining single-stranded RNA and oligos were then digested by adding 0.5uL of Buffer EB (Qiagen #19086), 0.5uL of thermolabile ExoI (NEB #M0568S), and 0.25uL of RNase A (ThermoFisher #EN0531), and incubating the samples at 37C for 30min, followed by heat inactivation at 85C for 5min and a hold step at 4C.

Up to 8 samples with different index sequences were then pooled, and cDNA was purified by performing a 1.4x left-sided SPRI selection (Beckman Coulter #B23317). Q5 UltraII Mastermix (NEB #M0544L) was then used along with partial Read1 and partial Read2 primers to amplify the cDNA for 7 cycles using a C1000 Touch Thermal Cycler (Bio-Rad #1851197), following the “CheapSeq PCR” parameters described below. Amplified cDNA was purified through a 0.7x left-sided SPRI selection (Beckman Coulter #B23317). Final library construction was performed by using Q5 UltraII Mastermix (NEB #M0544L) along with standard TruSeq indexing primers (NEB #E6440S) and amplifying for 8 cycles following the “CheapSeq PCR” parameters described below. Libraries were purified by performing a 0.4x-0.65x double-sided SPRI selection (Beckman Coulter #B23317). Purified libraries were assessed using Tapestation HSD1000 (Agilent #5067-5584 and #5067-5585). For oligo sequences, see **Supplementary Table 14**.

CheapSeq PCR:

1. 98C for 30s
2. X cycles of 98C for 10s, 65C for 75s
3. 65C for 5min
4. 4C forever

### Differentially Expressed Gene (DEG) Analysis of Bulk RNA-seq from CasRx-KD NPCs

Three distinct NPC samples were analyzed per condition. Individual KD conditions were compared to three non-targeting control (NTC) conditions, which were considered together as a set of 9 control samples.

Gene expression was quantified using alevin-fry with UMI deduplication (salmon-v1.9). A salmon index and transcript-to-gene map were built using Gencode v45. Counts from distinct transcripts that mapped to the same gene were summed. Only genes with a baseMean expression of >10 in NTC samples were considered for further analyses. Differential expression analysis was performed using DESeq2 (v1.42.0), with the model ∼Batch + sgRNA.

For KD confirmation, p-values were adjusted using Benjamini-Hochberg correction for multiple hypothesis testing for the set of 20 directly targeted genes. The raw p-value for SRCAP was derived from Droplet Digital PCR (see below), as SRCAP was filtered from RNA-seq analysis due to low expression. See **Supplementary Table 2** for details.

For transcriptome-wide analyses, p-values were adjusted using Benjamini-Hochberg correction for multiple hypothesis testing for the set of 11,168 analyzed genes (baseMean >10 in NTCs).

### Droplet Digital PCR (ddPCR)

To confirm KD of SRCAP, which was filtered from RNA-seq analysis due to low expression, the same RNA samples that had been used for RNA-seq (above) were assessed by ddPCR. RNA was first converted to cDNA using the SuperScript™ IV VILO™ Master Mix (ThermoFisher Scientific # 11756050). Next, ddPCR was performed by combining cDNA at an RNA-equivalent concentration of 0.25ng/uL (5ng per 20uL reaction), primers at a final concentration of 0.1uM, and QX200™ ddPCR™ EvaGreen Supermix (Bio-Rad # 1864033). The ddPCR reactions were performed using the QX200 AutoDG Droplet Digital PCR System (Bio-Rad). Expression levels were normalized using the expression of the housekeeping gene *HPRT1*. The results were analyzed using the model Expr ∼ Batch + type, where type was either NTC or the targeted gene (SRCAP). The resulting raw p-value was adjusted along with those for the other KD targets (from RNA-seq, above) using Benjamini-Hochberg correction for multiple hypothesis testing for the set of 20 directly targeted genes. See **Supplementary Table 2** for details.

### Primer Design

Primers (**Supplementary Table 15**) were designed using Primer-BLAST^84^, which assesses potential off-target interactions using BLAST^71^. Oligos were ordered through Azenta or IDT. Primer specificity was confirmed by Sanger sequencing of PCR products (Azenta).

### Auto-Western Blot

To analyze protein expression levels following CasRx KD, LiOn-based KD constructs (see “Molecular Cloning”) were nucleofected into 280-NPCs along with a piggyBac-transposase-expressing plasmid (see “Nucleofection” for details). Four days later, EGFP+ cells expressing the KD constructs were collected by FACS (BD FACSymphony™ S6). The cells were pelleted and flash-frozen in liquid nitrogen. Protein extraction and auto-western blot were performed by RayBiotech using the following antibodies: Vinculin (R&D systems MAB6896, clone 728526) diluted 1:25, and WAC (Abcam AB109486) diluted 1:50.

### Gene Sets

1. SFARI genes: all genes included in the SFARI gene database^14^ (4/3/25 release).
2. High confidence SFARI (HC-SFARI) genes: SFARI genes with a score of 1.
3. Intellectual disability (ID) genes: genes from the SysNDD database^85^ (accessed 8/23/25) that were:

a. associated with an ID term (HP:0001249, HP:0001256, HP:0002342, HP:0010864, HP:0002187)
b. in the “definitive” category
c. not SFARI genes

### Enrichment Analyses

Enrichment analyses were performed using one-sided Fisher’s exact tests (*fisher.test,* base R v4.3.1), and p-values were adjusted using Benjamini-Hochberg correction for multiple hypothesis testing (*p.adjust, method = “BH”,* base R v4.3.1). When displayed as bar plots, main bars indicate the odds ratio (OR) from a one-sided Fisher’s exact test; error bars indicate the lower bound of the 95% confidence interval on the OR (no upper bound is defined for a one-sided test).

### Knockdown of Fluorophores

Fluorophore-expressing constructs (see “Molecular Cloning”) were nucleofected into 280-iPSCs along with a piggyBac-transposase-expressing plasmid (see “Nucleofection” for details). iPSCs were expanded, and then cells expressing all three fluorophores were isolated by FACS using a Cytek Aurora CS spectral cell sorter. The sorted iPSCs were further expanded and then induced into NPCs (see “Cell Culture”). LiOn-based CasRx KD constructs were then nucleofected into these NPCs along with a piggyBac-transposase-expressing plasmid. Four days later, cells expressing all four fluorophores (with EGFP indicating expression of the KD constructs) were analyzed for fluorescence intensity of the reporters using the Cytek Aurora CS spectral cell sorter. Ordinary two-way ANOVA with Bonferroni’s multiple comparisons test was used to test for significant differences in the geometric mean fluorescence intensity (log_2_) across samples and reporters.

### Knockdown by shRNA

For each target, pSicoR-based constructs with three different shRNAs were pooled and then nucleofected into 280-NPCs (see “Nucleofection” for details, with modifications as described here). Since these constructs are not transposable, the plasmid expressing piggyBac transposase^73^ was omitted, and instead the amount of pSicoR-based plasmid was increased to 15ug total. The day after nucleofection, EGFP+ cells expressing the constructs were isolated by FACS (BD FACSymphony™ S6) and re-plated to allow recovery overnight. The following day, RNA was collected using Zymo DNA/RNA Shield. 3’ RNA-seq was performed by Plasmidsaurus (DGE RNA-Seq service).

### DEG Analysis of Bulk RNA-seq from shRNA-KD NPCs

Differential expression analysis was performed using the same analytical pipeline as our CasRx-KD experiments: DESeq2 (v1.42.0) was used to perform differential expression analysis with the model ∼targetGene, with differential expression calculated relative to NTC shRNAs (designed to target non-expressed transcripts luciferase and lacZ). For transcriptome-wide analyses, p-values were adjusted using Benjamini-Hochberg correction for multiple hypothesis testing for the set of 14,601 genes with baseMean >10 in NTC cells. Differentially expressed genes (DEGs) were genes that were differentially expressed with an adjusted p-value < 0.05.

### Analysis of Gene Expression Data from Deneault et al. (2018)

Gene expression counts from Deneault et al. (2018)^20^ were downloaded from GEO (GSE107878). Differential expression analysis was performed using the same analytical pipeline as our CasRx-KD experiments: DESeq2 (v1.42.0) was used to perform differential expression analysis with the model expression ∼ targeted gene. Differentially expressed genes (DEGs) were identified relative to control cells for both neurons and induced pluripotent stem cells. Only genes with a mean gene expression over 10 in control samples were considered as potential DEGs, and p-values were adjusted accordingly using a Benjamini-Hochberg correction. DEGs were identified as genes that were differentially expressed with an adjusted p-value < 0.05.

### Cross-Enrichment Analysis

We quantified the overlap between DEGs identified from: (1) our CasRx-KD experiments in NPCs, (2) our shRNA-KD experiments in NPCs, (3) KO iPSCs from Deneault et al. (2018)^20^, and (4) KO ExNs from Deneault et al. (2018)^20^. For each pairwise comparison of DEGs, enrichment analyses were performed using one-sided Fisher’s exact tests with a universe of genes that were expressed by both sample types being considered. P-values were adjusted using Benjamini-Hochberg correction for multiple hypothesis testing.

### Analysis of Expression Correlation with SFARI Genes

To determine whether a gene of interest was significantly co-expressed with SFARI genes in our NPC RNA-seq data, we calculated the median Pearson correlation between the gene of interest and each SFARI gene. To assess whether this degree of correlation was significant, we created an empirical null distribution by calculating the median correlation between the gene of interest and 1000 randomly selected gene sets of the same size as the SFARI gene set, using the universe of NPC-expressed genes (baseMean > 10 in NTCs). A one-sided empirical p-value was then calculated as the proportion of null values greater than or equal to the observed median correlation. Regulators with p-values < 0.05 were considered significantly correlated with SFARI genes.

### Identification of Consistently Downregulated DEGs

Consistently downregulated DEGs were defined as genes that exhibited: 1) statistically significant downregulation by at least three of the KDs, and 2) at least 80% concordance in the direction of significant differential expression across all KDs.

### Constraint Analysis

LOEUF scores (oe_lof_upper) were obtained from gnomad v2^86^ to evaluate genic constraint. For each gene, we calculated a per-gene LOEUF score by calculating the median of the LOEUF values of each gene’s associated transcripts. While LOEUF is a continuous metric, we utilized LOEUF < 0.35 as recommended to identify highly constrained genes.

### Additional Support for Candidate ASD Risk Genes

Candidate ASD Risk Genes (**Supplementary Table 8)** that were identified in any of the following major sequencing studies were included in **Supplementary Table 9**:

1. Trost et al. (2022)^2^
2. Zhou et al. (2022)^55^
3. Wang et al. (2022)^87^
4. Doan et al. (2019)^88^
5. Rodin et al. (2021)^89^
6. Wilfert et al. (2021)^90^

Additional references:

1. Alcantara et al. (2017)^91^
2. Marcelis et al. (2024)^92^
3. Ruiz-Reig et al. (2024)^93^
4. McGhee et al. (2025)^94^

### Nucleofection

Nucleofections were performed using the Neon Transfection System (ThermoFisher Scientific). For NPCs, 2M cells were nucleofected with 10ug of transposable plasmid and 2ug of a plasmid expressing piggyBac transposase^73^ using the 100uL tips. For iPSCs, 1M cells were nucleofected with 5ug of transposable plasmid and 1ug of a plasmid expressing piggyBac transposase^73^ using the 100uL tips. For small-scale nucleofections, 10uL tips were used and the cell number and plasmid amounts were reduced proportionally. Nucleofection of NPCs was performed using the following settings: voltage = 1600V, width = 10, pulses = 3. Nucleofection of iPSCs was performed using the following settings: voltage = 950V, width = 30, pulses = 2.

### NPC Proliferation Assays

LiOn-based KD constructs (see “Molecular Cloning”) were nucleofected into 280-NPCs along with a piggyBac-transposase-expressing plasmid (see “Nucleofection” for details). Four days later, the cells were treated for 20 minutes with dye from the CYTOTRACK™ Red 628/643 Cell Proliferation Assay Kit (Bio-Rad #1351205) diluted 1:500 in media. The cells were then rinsed with PBS (ThermoFisher Scientific # 10010031) and passaged, with 1/5 of each sample plated into a well of 4 separate plates to be used for the subsequent 4 timepoints. The remaining cells were analyzed by flow cytometry (BD FACSDiva software v8.0.3) to assess the initial CytoTrack intensity on day 0 (d0). For the next 4 days, 1 well of each sample was dissociated each day and analyzed by flow cytometry. The CytoTrack intensity values (negative log2) from the EGFP+ cells expressing the KD construct were normalized to the EGFP-cells from the same well to control for well-to-well variability in proliferation rates. They were further normalized to their initial CytoTrack intensity from d0 to control for variability in dye incorporation. The results for each KD were displayed relative to the mean value from the NTC samples, plotted as mean +/- standard error of the mean (SEM). Statistical analysis was performed using ordinary two-way ANOVA with Dunnett’s multiple comparisons test (GraphPad Prism). Full details regarding sample numbers and statistical results are available in **Supplementary Table 10**.

### Induction of FLICK Recombination

To induce expression of Cre and initiate recombination of the FLICK construct, cells/organoids were treated with 2ug/mL doxycycline (Sigma D9891-1G) for 24 hours.

### Assessment of FLICK KD Efficiency

To determine KD efficiency using FLICK constructs, three distinct samples were analyzed per condition. Control 280 iPSCs were nucleofected with constructs encoding the NTC1 control gRNA array or a gRNA array targeting the gene *ENSG00000261840*, along with a plasmid expressing piggyBac transposase^73^ to genomically integrate the FLICK constructs. To induce KD, the cells were treated with 2ug/mL doxycycline (Sigma D9891-1G) for 24 hours. Cells that had recombined the FLICK construct were then isolated through fluorescence-activated cell sorting (FACS) (BD FACSymphony™ S6) for EGFP+ cells, which were re-plated in triplicate and grown for an additional two days. RNA was then isolated using the Direct-zol RNA Microprep Kit (Zymo Research #R2062). To generate cDNA, 500ng of RNA was reverse transcribed using SuperScript™ IV VILO™ Master Mix (ThermoFisher Scientific #11756050). Quantitative PCR (qPCR) was performed by combining cDNA at an RNA-equivalent concentration of 1ng/uL (10ng per 10uL reaction), primers at a final concentration of 0.5uM, and PowerUp™ SYBR™ Green Master Mix for qPCR (ThermoFisher Scientific #A25742). The qPCR reactions were run in technical triplicate for each sample on the CFX384 Touch Real-Time PCR Detection System (BioRad #1855484) using the “fast cycling mode” conditions described in the Master Mix protocol. Expression of the KD target gene (*ENSG00000261840*) was assessed using the delta-delta Ct method, with expression first normalized to that of the housekeeping gene *HPRT1* for each sample, and then displayed relative to the average of the NTC control samples, plotted as mean +/- SEM. Statistical significance was assessed using a two-tailed t test (GraphPad Prism).

### Processing of Cerebral Organoids for Histological Analysis

Cerebral organoids were fixed in 4% paraformaldehyde (PFA), diluted in PBS from 16% PFA (Electron Microscopy Sciences # 15710), at 4°C overnight. Following fixation, cerebral organoids were rinsed with PBS and then equilibrated in 30% sucrose (w/v in PBS) at least overnight at 4°C. For large organoids, the 30% sucrose solution was replaced at least once following overnight incubation to ensure complete cryoprotection. For cryosectioning, cerebral organoids were frozen in Tissue-Tek® O.C.T. Compound (Sakura #4583) on dry ice. A Leica CM3050 S Cryostat was used to obtain 30um sections, which were collected on Fisherbrand™ Superfrost™ Plus Microscope Slides (Fisher Scientific #12-550-15) and stored at −80°C.

For immunohistochemical analysis, blocking buffer was prepared by combining 0.3M glycine (Sigma G7403-1KG), 1% bovine serum albumin (Sigma A4503-50G), 0.3% Triton X-100 (Sigma T8787-50ML), 10% normal goat serum (Fisher Scientific # NC9660079), and PBS. Slides were rinsed in PBS to remove O.C.T. and then incubated in blocking buffer for 1 hour at room temperature (RT). Blocking buffer was removed and replaced with fresh blocking buffer containing primary antibodies of interest (see below), and the slides were incubated in a slide moisture chamber (Avantor # 76278-848) at 4°C overnight. Slides were then rinsed with PBS three times for five minutes each. Secondary antibodies (see below) and the nuclear stain DAPI (ThermoFisher Scientific #62248) were diluted in blocking buffer (1:500 and 1:1000, respectively) and added to the slides for 30 minutes at RT, and the slides were protected from light from this point forward. The slides were then rinsed in PBS three times for five minutes each. Cover slips (Epredia #152455) were mounted using Fluoromount-G® (Fisher Scientific #OB100-01), which was allowed to dry at least overnight at RT. After drying, slides were stored at 4°C.

### Antibodies for Immunohistochemistry and Immunocytochemistry

1. PAX6 (BD Pharmingen 561462, clone O18-1330): diluted 1:300
2. Nestin (Millipore Sigma HPA026111-25UL): diluted 1:500
3. TBR2 (Millipore Sigma HPA028896-100UL): diluted 1:300
4. CTIP2 (Abcam ab18465, clone 25B6): diluted 1:300
5. EGFP (Aves GFP-1010): diluted 1:1000
6. Goat anti-Chicken IgY, Alexa Fluor 488 (ThermoFisher A-11039): diluted 1:500
7. Goat anti-Rabbit IgG, Alexa Fluor 488 (ThermoFisher A-11008): diluted 1:500
8. Goat anti-Rat IgG, Alexa Fluor 488 (ThermoFisher A-11006): diluted 1:500
9. Goat anti-Mouse IgG, Alexa Fluor 555 (ThermoFisher A-21424): diluted 1:500
10. Goat anti-Rat IgG, Alexa Fluor 647 (ThermoFisher A-21247): diluted 1:500
11. Goat anti-Rabbit IgG, Alexa Fluor 647 (ThermoFisher A-21245): diluted 1:500
12. Goat anti-Rabbit IgG, Alexa Fluor 750 (ThermoFisher A-21039): diluted 1:500

### Confocal Imaging and Processing

Cerebral organoid sections were imaged using a Zeiss LSM 980 confocal microscope. Standard image processing was performed using Adobe Photoshop.

### Cerebral Organoid scRNA-seq

For scRNA-seq, cells were collected from:

- 1 month: 10 organoids, pooled together and collected as a single batch
- 2 months: 14 organoids collected in two batches, with 7 organoids collected together on day 60 and an additional 7 organoids collected together on day 61
- acute KD: 8 organoids, pooled together and collected as a single batch

To collect cells, mosaic cerebral organoids were cut into pieces using spring scissors, rinsed twice with PBS to remove debris, and then dissociated using the Neural Tissue Dissociation Kit – Postnatal Neurons (Miltenyi Biotec # 130-094-802) using the enzyme mixes described for brain tissue. Gentle dissociation was performed by trituration with pipette tips of decreasing diameter, starting with wide orifice tips (Rainin # 30389218) and progressing to standard tips. Cells with recombined FLICK constructs were isolated through FACS (BD FACSymphony™ S6) for EGFP+ cells. Transcriptome-wide scRNA-seq libraries were prepared using Chromium Next GEM Single Cell 3ʹ Reagent Kits v3.1 (Dual Index) from 10X Genomics, following user guide CG000316 Rev A.

### Construction of FLICK Barcode Sequencing Libraries

Please see full details on protocols.io: https://dx.doi.org/10.17504/protocols.io.5qpvoj5pzg4o/v1. Briefly: for quantification of FLICK constructs from the initial pool of iPSCs used to generate mosaic organoids (d0), RNA was extracted from triplicate samples of 42k cells using the Direct-zol RNA Microprep Kit (Zymo Research #R2062). To generate cDNA, 50-250ng of RNA was reverse-transcribed using the SuperScript™ IV First-Strand Synthesis System (Invitrogen 18091050) along with an oligo(dT)VN primer containing UMIs. Sequencing libraries were then generated using Q5® Hot Start High-Fidelity 2X Master Mix (NEB #M0494L) and custom primers.

To generate FLICK barcode libraries from scRNA-seq libraries, 5uL of amplified and cleaned cDNA (produced by following the 10X User Guide CG000316 Rev A through the end of Step 2) was used. Barcode-containing cDNA was amplified for sequencing using Q5® Hot Start High-Fidelity 2X Master Mix (NEB #M0494L) and custom primers.

Libraries were cleaned using SPRIselect DNA Size Selection Reagent (Beckman Coulter # B23317) and assessed using TapeStation High Sensitivity D1000 reagents (Agilent #5067-5603 and #5067-5584).

### Alignment and Quantification of Organoid scRNA-seq Data

scRNA-seq data was aligned to the pre-built CellRanger (v.9.0.0) hg38 reference using *cellranger-count* in feature-barcode mode to perform simultaneous gene expression and guide abundance quantification. Introns were not included in gene expression quantification (--include-introns=false) to avoid quantifying partially degraded transcripts that had been targeted by CRISPR-CasRx. CRISPR guide sequencing library FASTQs for 2-month samples were down-sampled to 1.25% of reads using seqtk (v1.4) to match CRISPR library sequencing depth for one-month samples and to minimize background for guide assignment using the cellranger-CRISPR guide quantification pipeline. All other default parameters were used. CellRanger guide assignments were used to filter to only cells with exactly one assigned guide.

### Pre-Processing and Quality Control of Organoid scRNA-seq Data

Long-term and acute KD samples were processed separately but equivalently. CellBender (v0.3.2) was used to perform detection and filtering of ambient RNA contamination, while scrublet (v0.2.3) was used to perform automatic doublet detection and filtering. All other downstream analysis was performed with Seurat (v5.0.0). High-quality cells were retained according to the following quality-control metrics: 1) percent mitochondrial (percent.mt) gene expression < 20% and 2) 500 < number of expressed genes (nFeature_RNA) < 6000.

### Cell-Type Annotation of Organoid scRNA-seq Data

Cell-type annotation was first performed on only non-targeting control (NTC) cells. NTC cells were extracted to create a control-only Seurat object. This object subsequently underwent normalization (*NormalizeData*), variable feature identification (*FindVariableFeatures*), scaling (*ScaleData*; regressed covariates of percent.mt, percent_ribo, nCount_RNA, and nFeature_RNA), and dimensionality reduction (*RunPCA* and *RunUMAP*). Integration was performed using harmony (v1.2.0) across organoid batches and this integrated object was used for unsupervised clustering (*FindNeighbors* and *FindClusters*, resolution = 0.6 and 1.2). A small number of unsupervised clusters (<1000 cells) were filtered due to low cluster-wide QC metrics (i.e. exceptionally high or low nCount_RNA, high percent.mt). All subsequent 2-D visualization of this control-only Seurat object was performed using a UMAP representation of the integrated data. Cell type annotation was performed using canonical brain marker genes. A Seurat object was next constructed for all cells, following the same processing pipeline (e.g., normalization, scaling, dimensionality reduction, and integration) as described above. To perform cell-type annotation, label transfer was performed from the control-only Seurat object (*TransferData*) to the full Seurat object. Expression patterns of canonical brain marker genes were then used to refine and finalize these cell type annotations.

### Analysis of Relative Abundance per KD in Cerebral Organoids

The relative abundance of cells for each KD target was determined for each time point (d0, d30, d60, or d61). For d0, relative abundance was determined from bulk RNA-seq libraries (see “Construction of FLICK Barcode Sequencing Libraries”) using a custom pipeline (see **Code Availability**). For d30, d60, and d61, relative abundance was determined from scRNA-seq using the CRISPR guide assignments generated by CellRanger (see “Alignment and Quantification of Organoid scRNA-seq Data”). For each sample, results were normalized to the starting proportion at d0 and displayed relative to NTCs from the same timepoint. Results were plotted as mean +/- SEM. Statistical analysis was performed using repeated measures one-way ANOVA with Dunnett’s multiple comparisons test (GraphPad Prism), considering all long-term samples (d30, d60, and d61).

### Analysis of Relative Cell Type Proportions in Cerebral Organoids

The relative ExN/RG proportion was determined for each KD target at d30, d60, and d61. Results were normalized to the NTC sample from the same timepoint. Statistical analysis was performed using ordinary one-way ANOVA with Dunnett’s multiple comparisons test (GraphPad Prism), considering all long-term samples (d30, d60, and d61).

### Differential Expression Analysis of Cerebral Organoid scRNA-seq Data

Differential expression analysis was performed between perturbed cells per guide and cells that received NTCs, stratified by major cell types (radial glia, intermediate progenitor cells, excitatory neurons, or inhibitory neurons). Seurat *FindMarkers* was used to perform differential expression with logfc.threshold = 0, min.pct = 0.10, min.cells.group = 3 and otherwise all default parameters. Enrichment analyses of differentially expressed genes (DEGs) were performed using one-sided Fisher’s exact tests, and p-values were adjusted using Benjamini-Hochberg correction for multiple hypothesis testing. The universe was defined as all genes expressed in both comparisons of interest.

### Visualization of Shared DEGs Between Organoid Cell Types

Shared DEGs were visualized between organoid cell types (**Fig. 3c**) using the R package igraph (v2.1.4). The Fruchterman-Reingold layout algorithm (*layout_with_fr*) was used to determine final node placement.

### MAGIC Analyses

MAGIC^95^ (v1.1) was used to identify transcription factors and cofactors whose targets are enriched in the DEGs from individual perturbations. The strategy of “gene body (promoter to the end of the last exon) plus 1Kb flanking sequence either side of the gene body” was used. For NPC analyses, the universe was set as all genes with baseMean > 10 in NTC samples. For organoid analyses, the universe was set as all genes expressed in at least 10% of cells of that cell type (Seurat min.pct = 0.10; no log2 fold-change cutoff), as these were the genes assessed for differential expression.

### Re-Analysis of REST ChIP-Seq Data

Data was extracted from Table S2 from Rockowitz & Zheng (2015)^66^. Mouse genes were mapped to human genes using Orthology.eg.db (v3.20.0), org.Mm.eg.db (v3.20.0), and org.Hs.eg.db (v3.20.0). Enrichment analysis was performed using a one-sided Fisher’s exact test with all BrainSpan genes with normalized expression >= 1 as the universe (version “RNA-Seq Gencode v10 summarized to genes”; accessed 1/6/24; https://brainspan.org). P-values were adjusted using Benjamini-Hochberg correction for multiple hypothesis testing. Results were displayed as bar plots, with main bars indicating the odds ratio (OR) from a one-sided Fisher’s exact test and error bars indicating the lower bound of the 95% confidence interval on the OR (no upper bound is defined for a one-sided test).

### Gene Regulatory Network (GRN) Construction and Benchmarking

A GRN was constructed using the bioinformatic tool ARACNe-AP^96^. The input was the RNA-seq data from our 3 NTCs and the successful KD samples with at least 50 DEGs. We used curated lists of transcription factors^97^ and epigenetic regulators^98^ and selected genes expressed in NPCs (baseMean >10 in NTC samples) as the candidate genetic regulators. To create the GRN, we used default parameters and 100 bootstraps that were then aggregated together to form the final network. We then used VIPER^99^ (v1.36.0) to identify differentially active regulons following standard operating procedures. In brief, we identified regulons with significant differential expression between NTCs and KDs using a t-test, with a null distribution defined by sample permutations.

To benchmark the predictive power of the ARACNe-derived GRNs, we generated additional ARACNe networks after individually holding out perturbation samples for five target genes that were identified as regulators of at least 250 genes (range: 478-1219 genes) in their 2^nd^ degree regulons (i.e., their direct targets, which constitute their 1^st^ degree regulon, as well as the direct targets of those genes). For each held-out regulator, we assessed overlap using a one-sided Fisher’s exact test between 1) the predicted positively regulated genes within its 2^nd^ degree regulon, and 2) the set of downregulated DEGs resulting from perturbation of that same regulator. P-values were adjusted using Benjamini-Hochberg correction for multiple hypothesis testing. P-values across the five held-out perturbations were combined using Fisher’s method.

### PageRank Analysis of GRN

To identify central regulators within our GRN, we converted the GRN to a graph using the R package igraph (v2.1.4) and ran the PageRank algorithm^100^ using the igraph method *page_rank*. We then rank-ordered genes by their PageRank score and ran gene set enrichment analysis (GSEA)^101^ using the R package fgsea (v1.32.4), testing the following ontologies: SFARI genes, HC-SFARI genes, ID genes (exclusive of SFARI genes), and the 50 Molecular Signature Database Hallmark Pathways^102^ (obtained from https://www.gsea-msigdb.org/gsea/msigdb).

### Identification of Top Candidate Regulators for Driving Convergence in Autism

VIPER^99^ (v1.36.0) was used to identify GRN regulators whose regulons were significantly differentially active across the KDs compared to NTC controls. From these, we prioritized 70 regulators whose regulons were significantly enriched for both SFARI genes (one-sided Fisher’s exact test with Benjamini-Hochberg correction for multiple hypothesis testing) and high PageRank scores (gene set enrichment analysis with Benjamini-Hochberg correction for multiple hypothesis testing).

### Additional GRN Analyses

Mean normalized degree for the indicated set of regulators and their 1^st^ degree regulons (**Fig. 4d**) was calculated using the R package igraph (v2.1.4) by running the function *mean(degree(graph, normalized = TRUE))*. We then created an empirical null distribution of mean normalized degrees for each indicated set of regulators by creating 1M subnetworks of the same set size of randomly selected GRN regulators. Relative mean normalized degree was calculated as the observed mean normalized degree for a given set of regulators divided by the mean of the values for the corresponding empirical null distribution. A one-sided empirical p-value was then calculated as the proportion of null values greater than or equal to the observed mean normalized degree, with a detection limit of p = 1e-6.

To evaluate known or predicted protein-protein interactions between the top 70 regulators, we utilized the STRING database GUI^103^. For the universe, we used the full set genes that were considered as candidate genetic regulators in the GRN: transcription factors^97^ and epigenetic regulators^98^ that were expressed in NPCs (baseMean >10 in NTC samples). All other default parameters were used.

For clustering of our 70 regulators into modules, we used iterative walktrap clustering using the R package igraph (v2.1.4) by running the function *cluster_walktrap* with default parameters.

### Re-Analysis of CHD8 ChIP-Seq Data

For enrichment analysis of CHD8 targets, we used Supplementary Data 1 from Sugathan et al. (2014)^45^. Enrichment of the indicated gene sets was tested using one-sided Fisher’s exact tests, with the universe set to NPC-expressed genes (baseMean > 10 in NTCs). P-values were adjusted using Benjamini-Hochberg correction for multiple hypothesis testing. Results were displayed as a bar plot, with main bars indicating the odds ratio (OR) from a one-sided Fisher’s exact test and error bars indicating the lower bound of the 95% confidence interval on the OR (no upper bound is defined for a one-sided test).

### Identification of Female-Biased Genes

To identify genes that exhibit female-biased expression in the neurotypical brain, we combined DEG call sets from the following works:

1. Table S4 from Kissel et al. (2024)^68^: Identified as significantly female-biased in either the UCLA cohort, BrainVar cohort, or via meta-analysis.
2. Oliva et al. (2020)^104^ (https://storage.cloud.google.com/adult-gtex/bulk-qtl/v8/sb-eqtl/GTEx_Analysis_v8_sbgenes.tar.gz): Identified as significantly female-biased in either BRNCTXA or BRNCTXB.
3. Table S4 from Naqvi et al. (2019)^105^: Identified as significantly female-biased in the brains of all assessed species (human, mice, rats, dogs, and macaques).
4. Table S13 from DeCasien et al. (2026)^62^: identified as ubiquitously female-biased.

### Analysis of Single-Nucleus RNA-Sequencing (snRNA-seq) from Autistic Males

Processed snRNA-seq data from the human frontal cortex was obtained from Dias et al. (2024)^47^. Sex of individual samples was verified based on expression of *XIST* and Y-linked genes. Only a small number of female samples (one female control, one sample from a female with idiopathic autism, and one sample from a female with 15q-duplication-associated autism) were included in this cohort due to the known male bias of autism. Thus, we focused all downstream analysis on the 17 male samples included within this cohort. Differential expression analysis was performed using the Seurat function *FindMarkers* on a cell-type-specific basis between (a) male controls and just samples from males with idiopathic autism; (b) male controls and just samples from males with 15q-duplication-associated autism; and (c) male controls and all samples from males with autism, either idiopathic or 15q-duplication-associated. To control for donor effects, *FindMarkers* was run using MAST^106^. Significant genes were defined as those with an absolute log-2 fold-change > 0.1 and a transcriptome-wide adjusted p-value < 0.05.

### Plots of the Sex-Differential Expression of ZFX

Data from the following sources were used:

1. Benoit-Pilven et al. (2025)^67^
2. Kissel et al. (2024)^68^
3. Fass et al. (2024)^69^
4. DeCasien et al. (2026)^62^

### Re-Analysis of ZFX Targets from Rhie et al. (2018) and San Roman et al. (2024)

ZFX ChIP-seq targets were obtained from Table S3E from Rhie et al. (2018)^49^. ZFX targets were also obtained from Table S10 from San Roman et al. (2024)^48^, which previously performed integrative analysis of ZFX ChIP-seq data (ENCODE) and *ZFX* KD RNA-seq data from Rhie et al. (2018)^49^. Enrichment was analyzed using one-sided Fisher’s exact tests and p-values were adjusted using Benjamini-Hochberg correction for multiple hypothesis testing. For analysis of common ZFX targets from Rhie et al. (2018)^49^, the universe was defined as genes expressed in both of the cell lines used for ChIP-seq for which RNA-seq was also available (C4-2B and MCF-7). For analysis of ZFX targets from San Roman et al. (2024)^48^, the universe was defined as genes expressed in the corresponding cell type from Rhie et al. (2018)^49^. Results were displayed as a bar plot, with main bars indicating the odds ratio (OR) from a one-sided Fisher’s exact test and error bars indicating the lower bound of the 95% confidence interval on the OR (no upper bound is defined for a one-sided test).

### Software Summary

Software used in this manuscript included R (v4.3.1) and the R packages DESeq2 (v1.42.0), Seurat (v5.0.0), igraph (v2.1.4), scrublet (v0.2.3), harmony (v1.2.0), Orthology.eg.db (v3.20.0), org.Mm.eg.db (v3.20.0), org.Hs.eg.db (v3.20.0), VIPER (v1.36.0), and fgsea (v1.32.4), as well as CellRanger (v9.0.0), CellBender (v0.3.2), MAGIC (v1.1), salmon (v1.9), seqtk (v1.4), and BD FACSDiva software v8.0.3.

## Supporting information

Guide to Supplementary Information

Source Data for Fig. 1

Source Data for Fig. 4

Source Data for Fig. 5

Source Data for Ext. Data Fig. 4

Source Data for Ext. Data Fig. 7

Source Data for Ext. Data Fig. 8

## ACKNOWLEDGEMENTS

We thank members of the Walsh Lab for helpful discussions. We are also grateful to Dr. Norma K. Hylton and Evan M. Bushinsky of the Walsh Lab for assistance with cerebral organoid culture, Dr. August Yue Huang for technical feedback regarding statistical analyses, and Dr. Adrianna K. San Roman and Dr. Marla E Tharp of the Page Lab for their insights regarding *ZFX*.

## AUTHOR CONTRIBUTIONS

R.E.A. and M.T. conceived the experiments and led the analyses, with supervision from C.A.W. M.T. led all bioinformatic analyses. R.E.A. designed all plasmid constructs with support from T.S. and S.L. Molecular cloning and lentivirus production were performed by R.E.A., T.S., and S.L. The CasRx knock-in iPSC line was generated by J.H.T.S. Cell culture was performed by R.E.A., T.S., S.L., and N.M., with contributions from S.Z., G.E., and J.H. Bulk RNA-seq libraries were prepared by R.E.A. and S.L. Analysis of bulk RNA-seq results was performed by M.T., with code contributions from R.N.D. NPC proliferation assays were performed by R.E.A., S.L., S.Z., G.E., and J.H. Cerebral organoids were cultured by R.E.A., T.S., and N.M., with advice and support from X.Q. Cerebral organoid scRNA-seq libraries were prepared by R.E.A. and N.M. Analysis of scRNA-seq libraries was performed by M.T. Cryosectioning and immunohistochemical labeling of cerebral organoids was performed by R.E.A. and N.M. Confocal imaging and histological analyses of cerebral organoids were performed by D.E.-A. D.C.P. contributed to the interpretation of *ZFX* as a contributor to the female protective effect in ASD. R.E.A., M.T., and C.A.W. wrote the manuscript with contributions from all of the authors. All of the authors read and approved the final manuscript.

## ETHICS STATEMENT

All human cell culture work was performed under protocols approved by Boston Children’s Hospital IRB #S07-02-0087.

## DATA AND CODE AVAILABILITY

Data Availability: All raw data and processed count matrices have been provided on the Gene Expression Omnibus (GEO GSE332574: https://www.ncbi.nlm.nih.gov/geo/query/acc.cgi?acc=GSE332574).

Differential gene expression results can also be found in **Supplementary Tables 16-65**. This study also made use of the following publicly available datasets and websites, and copies of these datasets are also available via the Github for this manuscript (https://github.com/MayaTalukdar/convergence_in_autism_ms).

- GENCODE v45 (reference transcriptome annotation): https://www.gencodegenes.org
- SFARI Gene database (4/3/25 release): https://gene.sfari.org
- SysNDD database (accessed 8/23/25): https://sysndd.dbmr.unibe.ch
- RNA-sequencing expression data from CRISPR-edited iPSC-derived neurons and iPSCs (Deneault et al., 2018): GEO accession GSE107878 (https://www.ncbi.nlm.nih.gov/geo/query/acc.cgi?acc=GSE107878)
- gnomAD v2 (LOEUF/oe_lof_upper constraint scores), Table S11 (Karczewski et al., 2020): https://doi.org/10.1038/s41586-020-2308-7
- ASD sequencing study, Table S5G (Trost et al., 2022): https://doi.org/10.1016/j.cell.2022.10.009
- ASD sequencing study, Tables S2 and S9 (Zhou et al., 2022): https://doi.org/10.1038/s41588-022-01148-2
- ASD sequencing study, Table S6 (Wang et al., 2022): https://doi.org/10.1073/pnas.2203491119
- ASD sequencing study, Table S9 (Doan et al., 2019): https://doi.org/10.1038/s41588-019-0433-8
- ASD sequencing study, Table S6 (Rodin et al., 2021): https://doi.org/10.1038/s41593-020-00765-6
- ASD sequencing study, Table S5 (Wilfert et al., 2021): https://doi.org/10.1038/s41588-021-00899-8
- ENCODE ChIP-seq compendium, accessed via MAGIC (Roopra, 2020): https://www.encodeproject.org
- Human and mouse REST/NRSF ChIP-seq binding sites, Table S2 (Rockowitz & Zheng, 2015): https://doi.org/10.1093/nar/gkv514
- BrainSpan RNA-Seq data, “Gencode v10 summarized to genes” (https://www.brainspan.org; accessed 1/6/24)
- MSigDB Hallmark gene set collection: https://www.gsea-msigdb.org/gsea/msigdb
- STRING protein-protein interaction database: https://string-db.org
- CHD8 ChIP-seq target genes, Table S1 (Sugathan et al., 2014): https://doi.org/10.1073/pnas.1405266111
- Sex-biased expression data, Table S4 (Kissel et al., 2024): https://doi.org/10.1016/j.bpsgos.2024.100321
- Sex-biased expression data (Oliva et al., 2020): https://storage.cloud.google.com/adult-gtex/bulk-qtl/v8/sb-eqtl/GTEx_Analysis_v8_sbgenes.tar.gz
- Sex-biased expression data, Table S4 (Naqvi et al., 2019): https://doi.org/10.1126/science.aaw7317
- Sex-biased expression data, Table S13 (DeCasien et al., 2026): https://doi.org/10.1126/science.aea9063
- Sex-biased expression data, Table S2 (Benoit-Pilven et al., 2025): https://doi.org/10.1016/j.xgen.2025.100890
- Sex-biased expression data, Table S1-S2 (Fass et al., 2024): https://doi.org/10.1186/s13293-024-00622-2
- Single-nucleus RNA-sequencing data from frontal cortex of autistic and control males (Dias, Mo et al., 2024): https://doi.org/10.1016/j.ajhg.2024.07.002
- ZFX ChIP-seq target genes, Table S3E (Rhie et al., 2018): https://doi.org/10.1101/gr.228809.117
- Integrated ZFX target gene analysis, Table S10 (San Roman et al., 2024): https://doi.org/10.1016/j.xgen.2023.100462

Code Availability: Scripts, analyses, and reference files used for this study are accessible in a GitHub repository (https://github.com/MayaTalukdar/convergence_in_autism_ms).

---

**Extended Data Figure 1.**
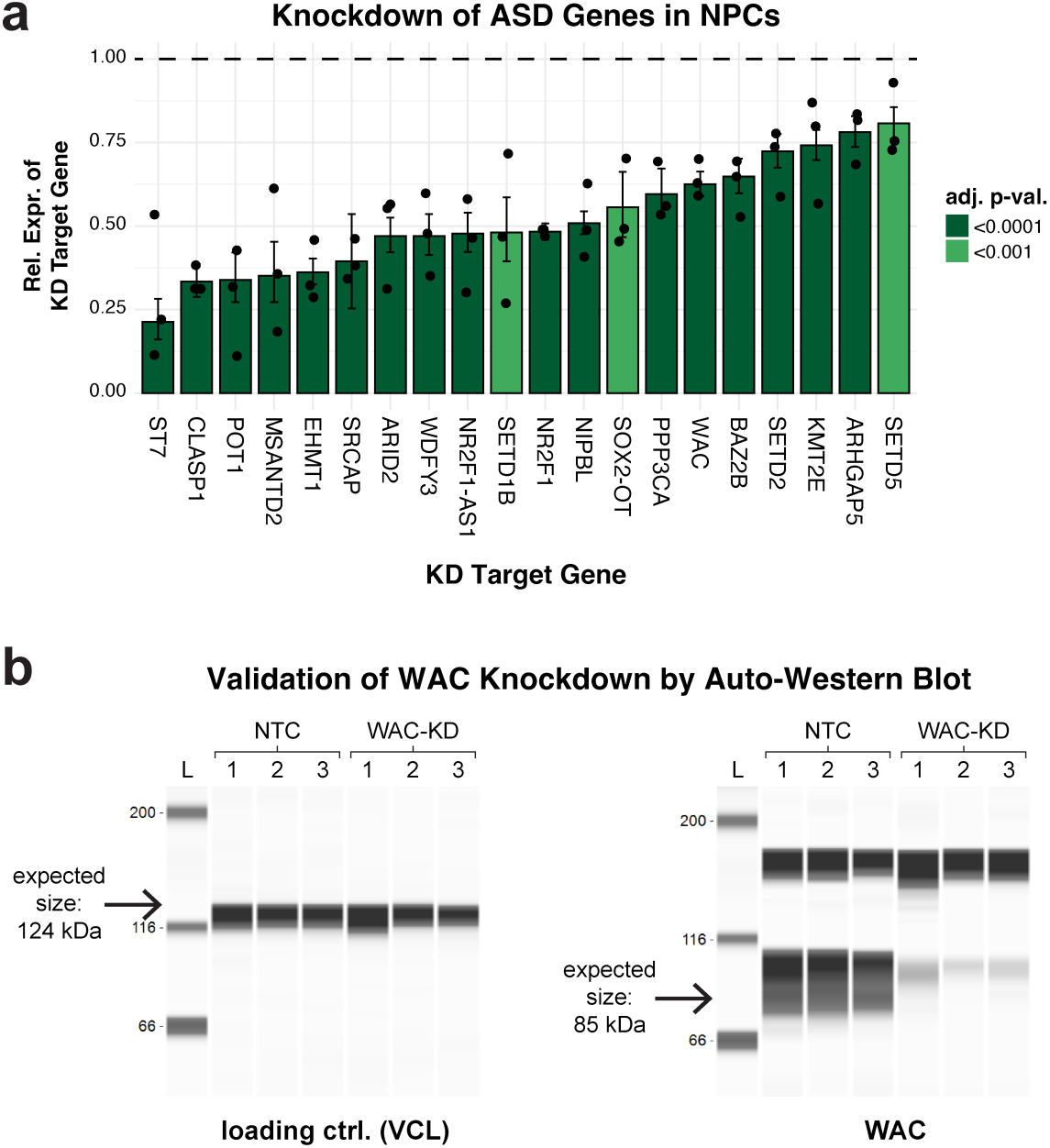
CasRx-based knockdown of ASD genes in NPCs. **a**, Relative expression of the targeted gene within each KD sample compared to NTC samples. Error bars: mean +/- 95% confidence interval. n = three biologically independent samples per condition. Statistically significant DEGs were identified using DESeq2 (Wald test) with Benjamini-Hochberg correction for multiple hypothesis testing. **b**, Analysis of protein levels by auto-western blot from NTC and WAC-KD NPCs, from three biologically independent samples per condition.

**Extended Data Figure 2.**
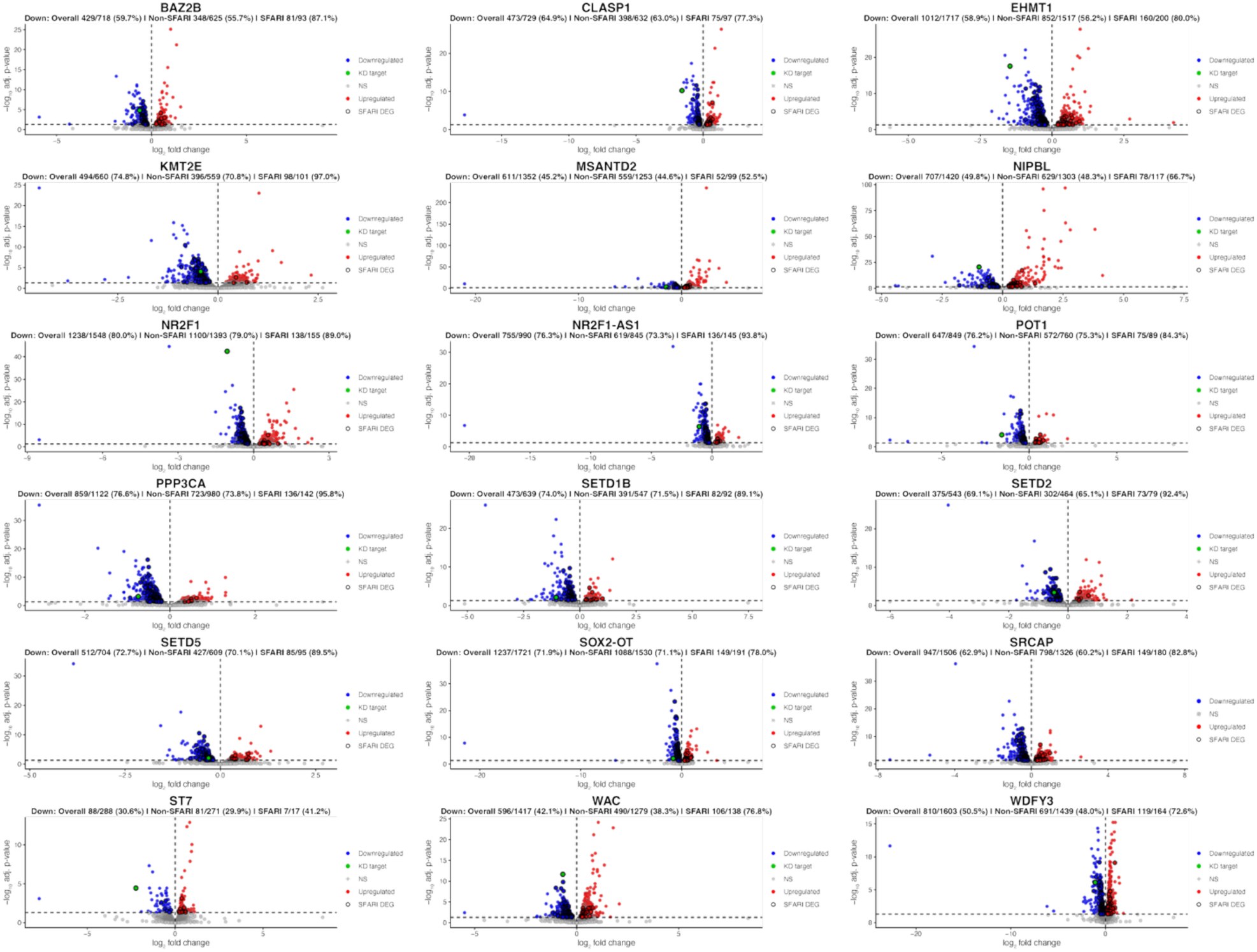
Differential gene expression upon CasRx-based KD of ASD genes in NPCs. Volcano plots depicting transcriptome-wide effects on gene expression following CasRx-based KD of ASD genes. Statistically significant DEGs were identified using DESeq2 (Wald test) with Benjamini-Hochberg correction for multiple hypothesis testing. Statistically significantly downregulated DEGs are shown in blue, while statistically significantly upregulated DEGs are shown in red. The gene targeted for KD (listed above each plot) is highlighted in green. SFARI DEGs are indicated by a thick border.

**Extended Data Figure 3.**
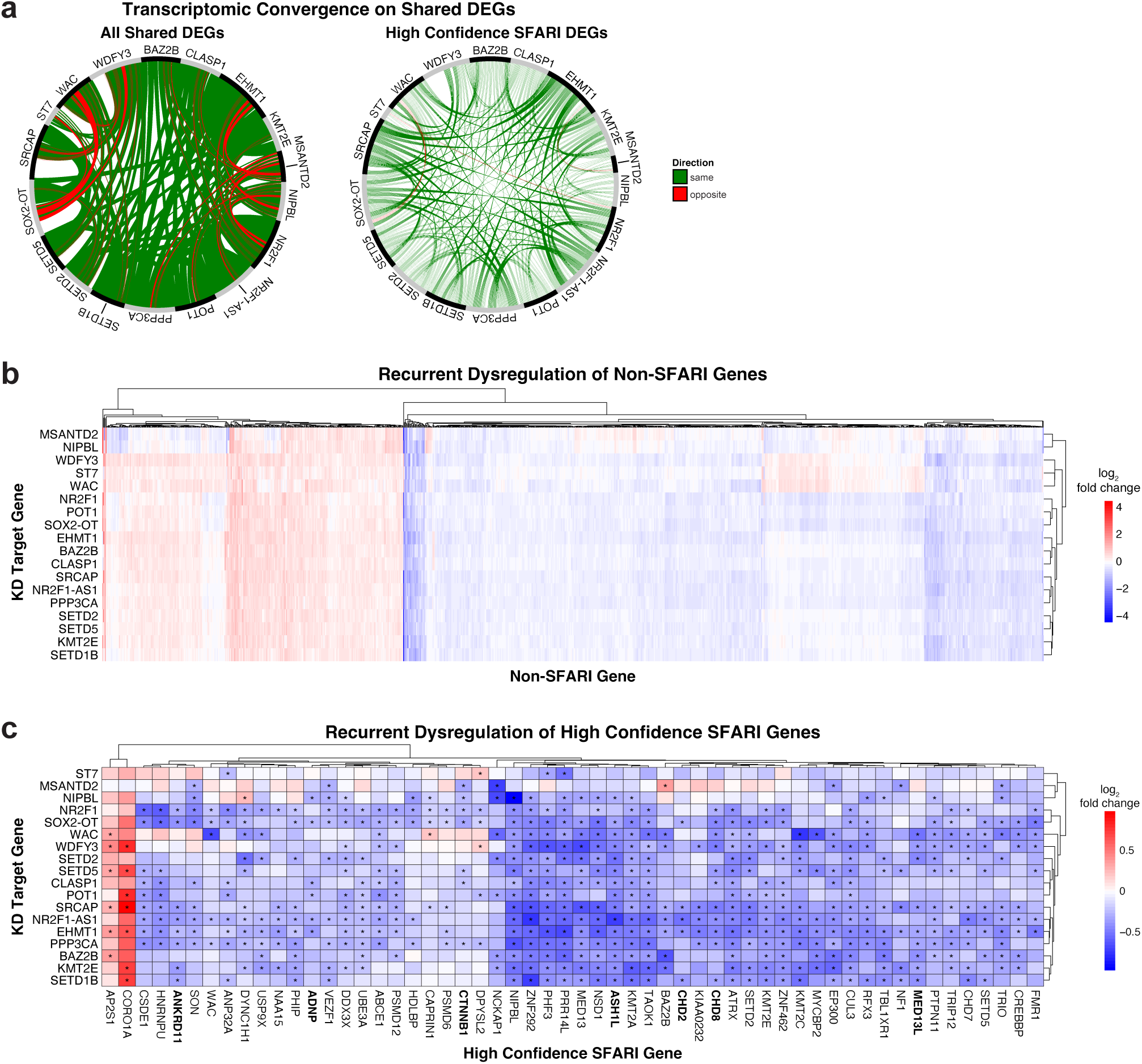
Transcriptomic convergence on shared differentially expressed genes. **a**, Circos plots depicting DEGs shared between different KD conditions. For direct comparison, the Circos plot for high confidence SFARI genes (shown in Fig. 1d) is also shown here. **b,** Heatmap showing the relative expression of non-SFARI genes that are DEGs in at least 5 KD conditions. **c,** Heatmap showing the relative expression of high confidence SFARI genes that are DEGs in at least 5 KD conditions. Statistically significant DEGs were identified using DESeq2 (Wald test) with Benjamini-Hochberg correction for multiple hypothesis testing, and are indicated with *. Genes that were identified amongst the 26 strongest ASD risk genes in Satterstrom et al. (2020)^1^ are bolded.

**Extended Data Figure 4.**
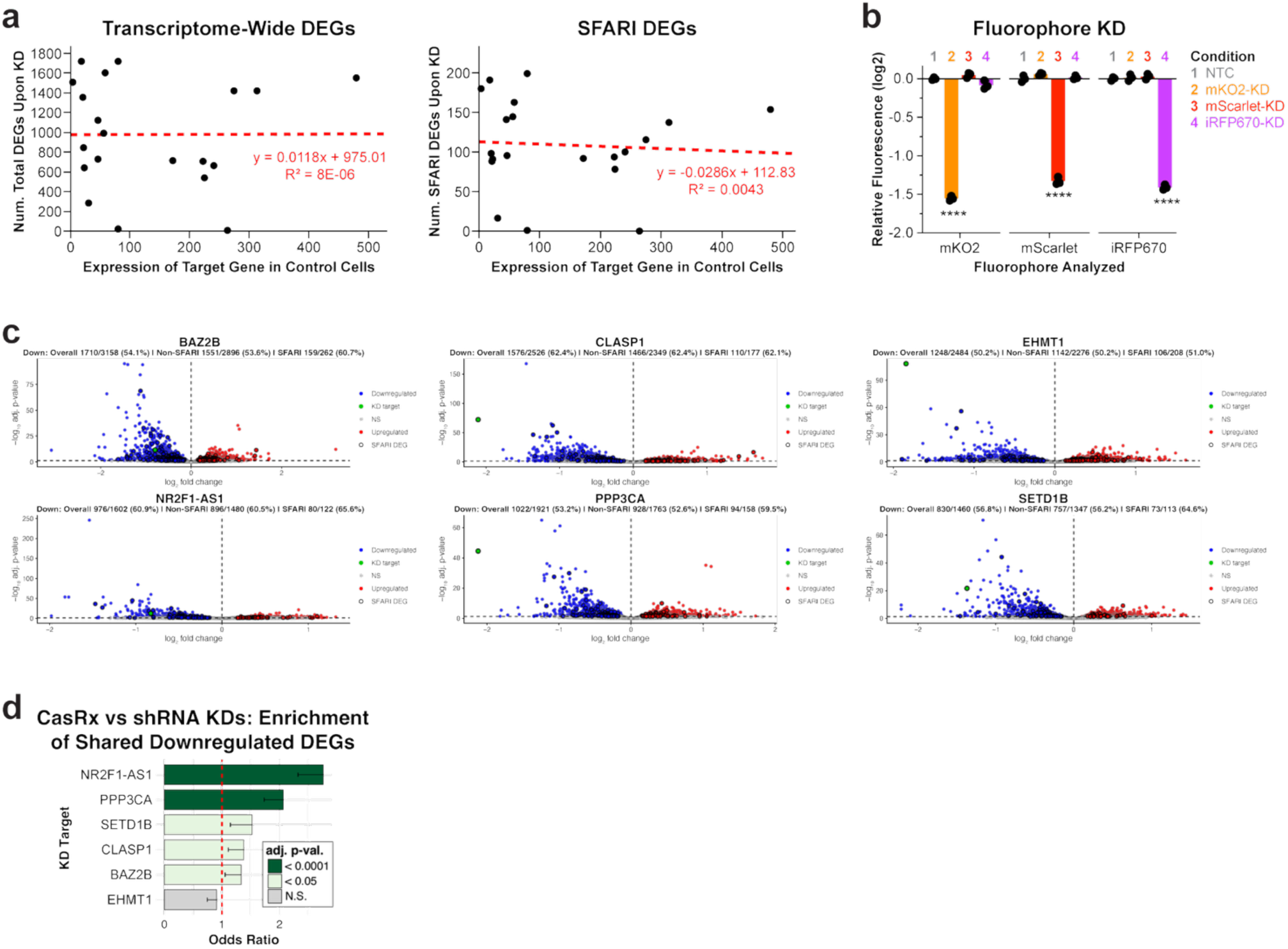
Shared DEGs are not dependent on CasRx-based KD. **a**, Scatter plots depicting the number of total DEGs (left plot) or SFARI DEGs (right plot) on the y-axis and the expression of the KD target gene on the x-axis. Linear regression is shown in red. **b,** Flow cytometry analysis of fluorescence intensity in NPCs expressing the fluorophores mKO2, mScarlet, and iRFP670. CasRx was used to individually KD each of the fluorophores. Results depicted relative to the NTC control average. **** adj. p = 1.44e-25 for mKO2, 5.74e-24 for mScarlet, and 1.45e-24 for iRFP670 (ordinary two-way ANOVA with Bonferroni’s multiple comparisons test). n = three biologically independent samples per condition. **c,** Volcano plots depicting transcriptome-wide effects on gene expression following shRNA-based KD of ASD genes. Statistically significant DEGs were identified using DESeq2 (Wald test) with Benjamini-Hochberg correction for multiple hypothesis testing. Statistically significantly downregulated DEGs are shown in blue, while statistically significantly upregulated DEGs are shown in red. The gene targeted for KD (listed above each plot) is highlighted in green. SFARI DEGs are indicated by a thick border. **d,** Bar plot depicting enrichment of the same downregulated DEGs following KD of the same target gene by CasRx vs shRNA. Bars indicate the odds ratio (OR) from a one-sided Fisher’s exact test; error bars indicate the lower bound of the 95% confidence interval on the OR (no upper bound is defined for a one-sided test). p-values were adjusted using Benjamini-Hochberg correction for multiple hypothesis testing.

**Extended Data Figure 5.**
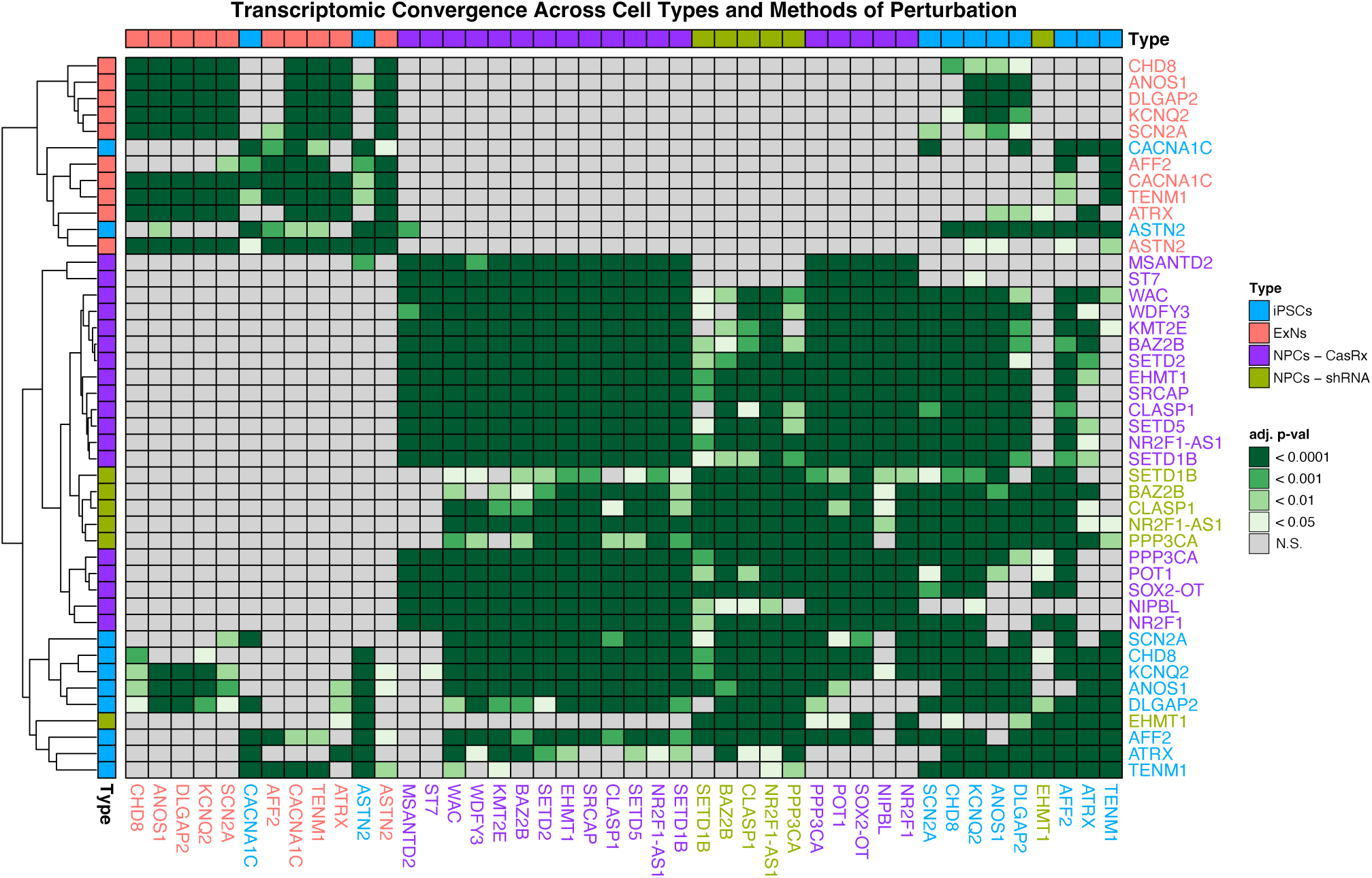
Transcriptomic convergence occurs across cell types and experimental contexts. Analysis of transcriptomic convergence across multiple cell types and experimental contexts. Statistically significant cross-enrichment of downregulated DEGs is shown in green (one-sided Fisher’s exact tests with Benjamini-Hochberg correction for multiple hypothesis testing). Data from iPSCs and ExNs were re-analyzed from Deneault et al. (2018)^20^.

**Extended Data Figure 6.**
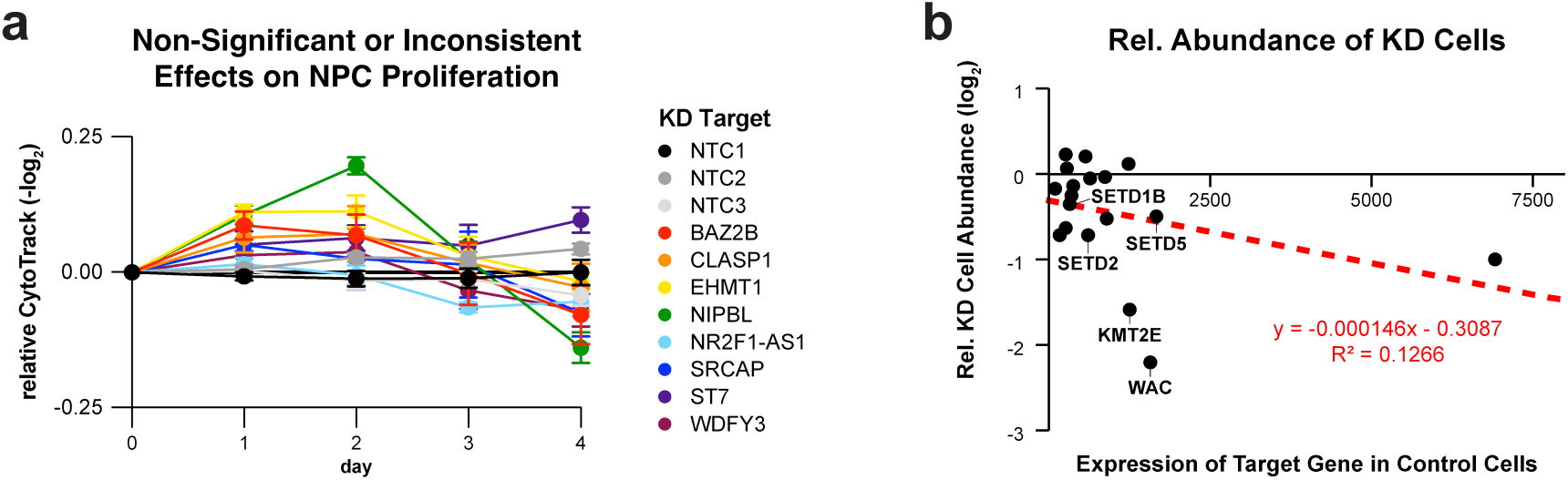
Effects on proliferation and cell abundance following ASD gene KD. **a**, Relative CytoTrack dye intensity (-log_2_) over time. Only KD conditions that exhibited non-significant or inconsistent effects are displayed. Plot depicts mean +/- standard error of the mean (SEM). Statistical significance was determined using ordinary two-way ANOVA with Dunnett’s multiple comparisons test. At least four biologically independent samples per condition were analyzed for each timepoint. See also Fig. 1f, and see **Supplementary Table 10** for full statistical results. **b,** Scatter plot depicting the relative abundance of KD cells on the y-axis and the expression of the KD target gene on the x-axis. Linear regression is shown in red. Genes that have been implicated in microcephaly and whose KD resulted in decreased proliferation (Fig. 1f) are highlighted.

**Extended Data Figure 7.**
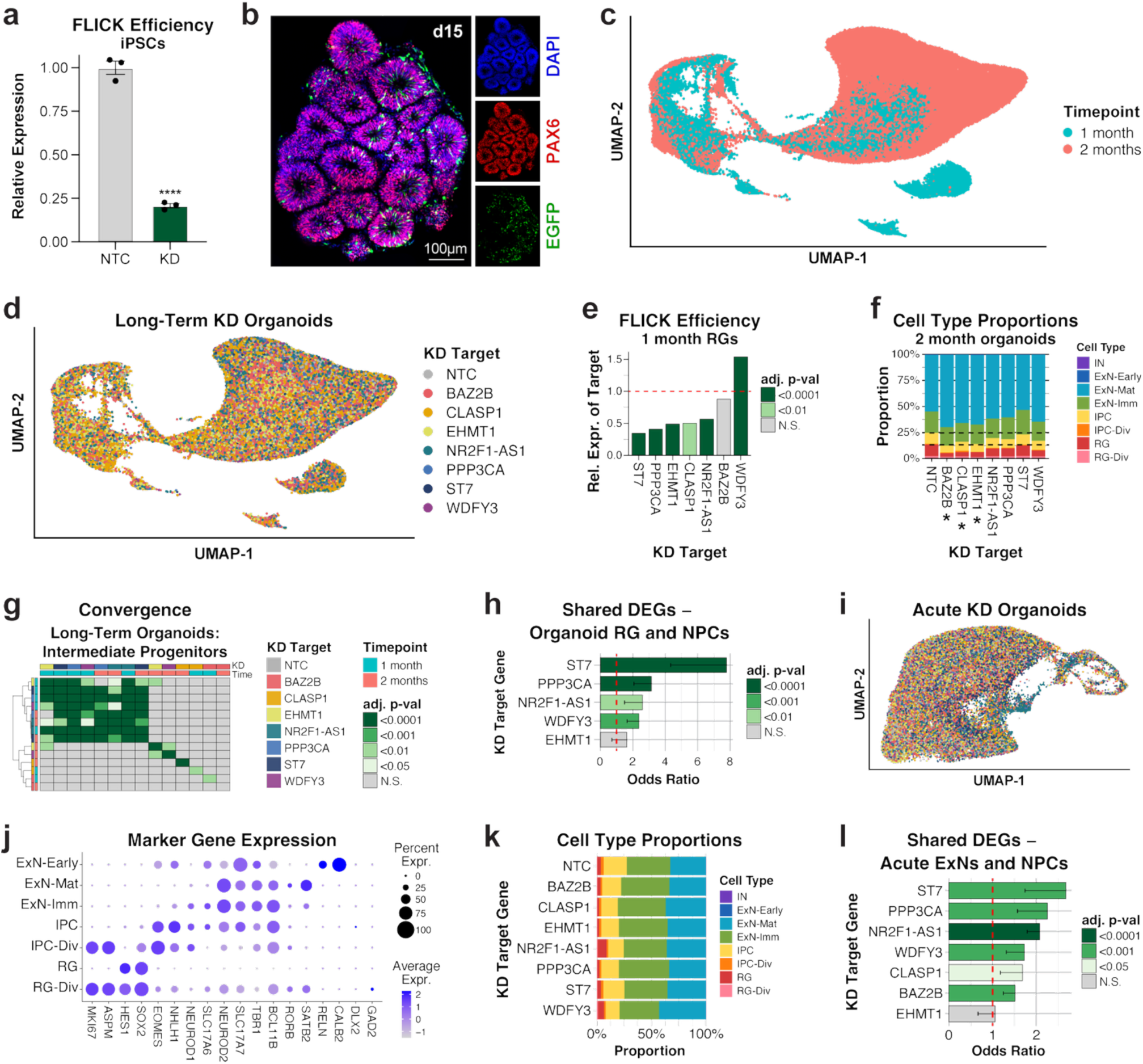
KD of ASD genes in mosaic cerebral organoids reveals transcriptomic convergence. **a**, KD efficiency of FLICK construct in iPSCs. Normalized expression of the KD target gene *ENSG00000261840* shown relative to NTC samples, as mean +/- SEM. **** p = 3.87e-5, two-tailed t test. n = three biologically independent samples per condition. **b,** Immunohistochemistry of a FLICK organoid at day 15, representative of three independent analyses with similar results. **c,** UMAP of organoid scRNA-seq colored by timepoint. **d,** UMAP of organoid scRNA-seq data colored by targeted gene. **e,** FLICK efficiency in RGs as quantified by scRNA-seq from one-month organoids. Statistically significant DEGs were identified using Seurat (non-parametric wilcoxon rank test) with Bonferroni correction for multiple hypothesis testing. We note that *BAZ2B* quantification may be impacted by low cell number, and *WDFY3* capture in scRNA-seq may be impacted by the length (>14kb) of the transcript. **f,** Cell type composition per KD target in two-month organoids. Dotted lines indicate the NTC proportions of major cell types. Samples for which the ExN/RG proportion was significantly different than NTC are indicated with * (adj. p = 0.021 for BAZ2B, 0.018 for CLASP1, and 0.012 for EHMT1), ordinary one-way ANOVA with Dunnett’s multiple comparisons test, considering all long-term samples from one month (see Fig. 2f) and two months (shown here). **g,** Cross-enrichment of DEGs from IPCs (one-sided Fisher’s exact tests with Benjamini-Hochberg correction for multiple hypothesis testing). **h,** Bar plot depicting cross-enrichment of DEGs from RGs from one-month organoids versus DEGs from previous NPC cultures upon KD of the same target gene. Only perturbations with at least 20 DEGs were tested for cross-enrichment. Bars indicate the odds ratio (OR) from a one-sided Fisher’s exact test; error bars indicate the lower bound of the 95% confidence interval on the OR (no upper bound is defined for a one-sided test). p-values were adjusted using Benjamini-Hochberg correction for multiple hypothesis testing. **i,** UMAP of scRNA-seq data from acute KD organoids, colored by KD target gene as in panel d. **j,** Expression of marker genes used for identifying cell types. **k,** Cell type composition across different perturbations in acute KD organoids. **l,** Bar plot depicting cross-enrichment of DEGs from excitatory neurons (ExNs) from acute KD mosaic organoids versus DEGs from previous NPC cultures upon KD of the same target gene. Bars indicate the odds ratio (OR) from a one-sided Fisher’s exact test; error bars indicate the lower bound of the 95% confidence interval on the OR (no upper bound is defined for a one-sided test). p-values were adjusted using Benjamini-Hochberg correction for multiple hypothesis testing.

**Extended Data Figure 8.**
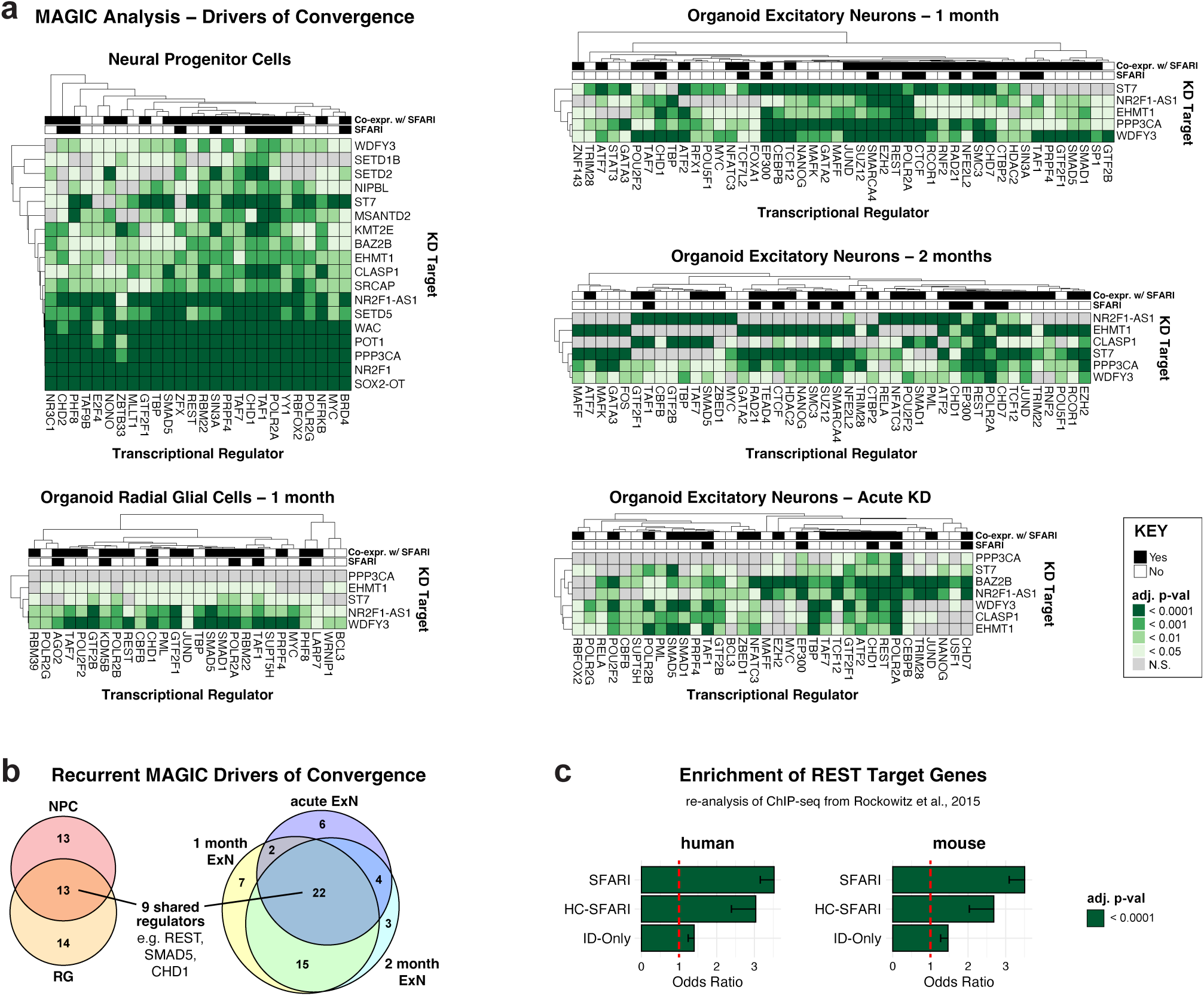
Key transcription factors, including REST, target recurrently dysregulated genes. **a**, MAGIC analysis (Kolmogorov-Smirnov test with Benjamini-Hochberg correction for multiple hypothesis testing) identifying transcriptional regulators whose targets were enriched within the DEGs from different cell types following KD of ASD genes. For each population, MAGIC regulators were scored by the number of ASD gene KDs that implicated the regulator. The 25 top scoring regulators are included here, along with any additional regulators with a tied score. **b,** Quantitative Euler diagrams depicting the overlapping MAGIC transcriptional regulators identified in different conditions. **c,** Bar plot depicting enrichment of SFARI, HC-SFARI, or ID-only genes within REST targets from human and mouse embryonic stem cells. REST target data from Rockowitz & Zheng (2015)^66^. Bars indicate the odds ratio (OR) from a one-sided Fisher’s exact test; error bars indicate the lower bound of the 95% confidence interval on the OR (no upper bound is defined for a one-sided test). p-values were adjusted using Benjamini-Hochberg correction for multiple hypothesis testing.

**Extended Data Figure 9.**
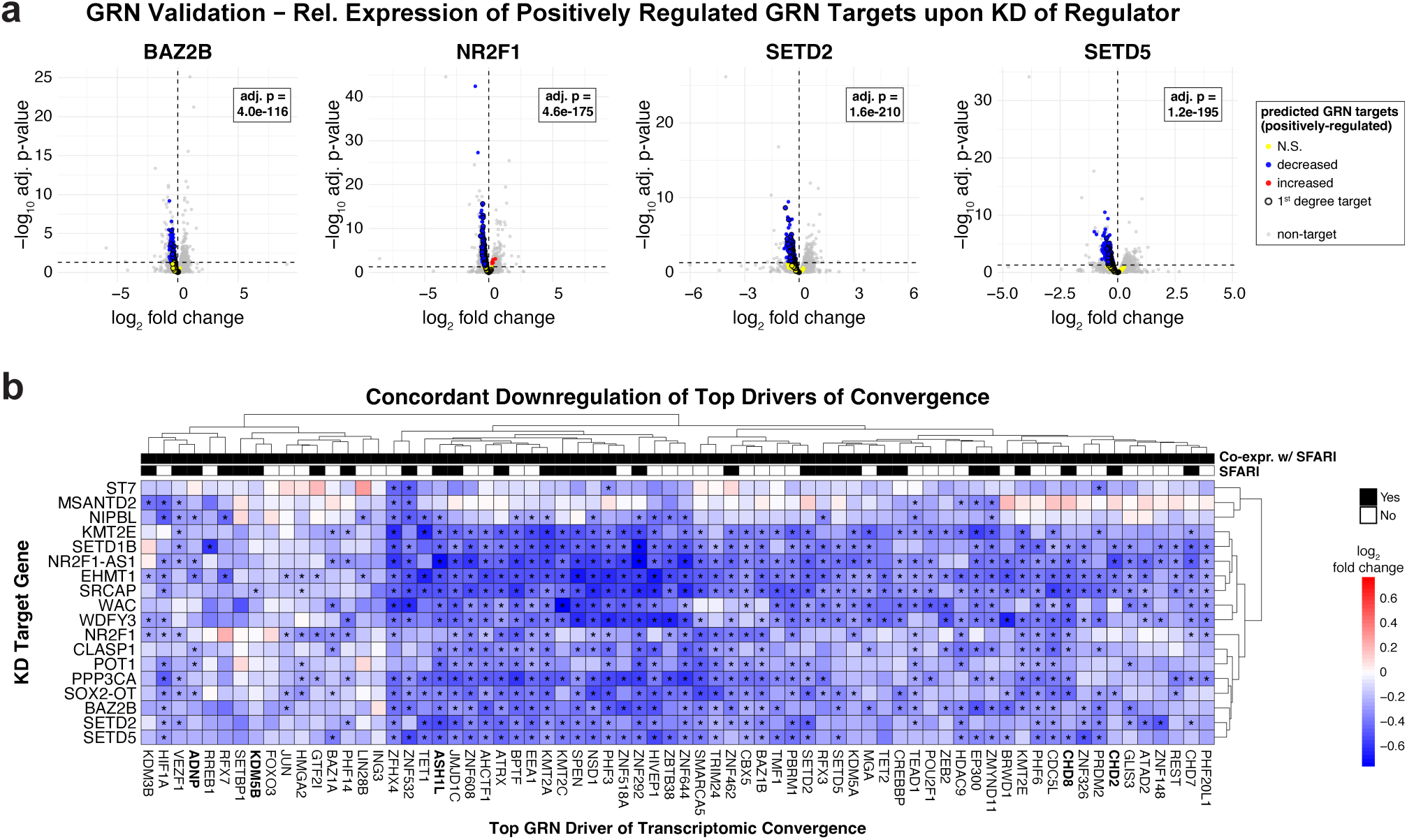
Top drivers of convergence identified through reconstruction of an NPC gene regulatory network. **a**, Volcano plots depicting differential expression upon KD of the regulator shown above the plot. Genes predicted by GRN analysis to be positively regulated by the indicated regulator are highlighted, with statistically significantly decreased DEGs shown in blue, increased DEGs shown in red, and targets with no significant change shown in yellow. Statistically significant DEGs were identified using DESeq2 (Wald test) with Benjamini-Hochberg correction for multiple hypothesis testing. Statistically significant enrichment of 2nd degree regulon genes within the downregulated DEGs is indicated by the adjusted p-value (one-sided Fisher’s exact tests with Benjamini-Hochberg correction for multiple hypothesis testing). **b,** Heatmap showing the relative expression of the top GRN drivers of convergence across the different ASD gene KD samples. Statistically significant DEGs were identified using DESeq2 (Wald test) with Benjamini-Hochberg correction for multiple hypothesis testing, and are indicated with *. Genes that were identified amongst the 26 strongest ASD risk genes in Satterstrom et al. (2020)^1^ are bolded.

**Extended Data Figure 10.**
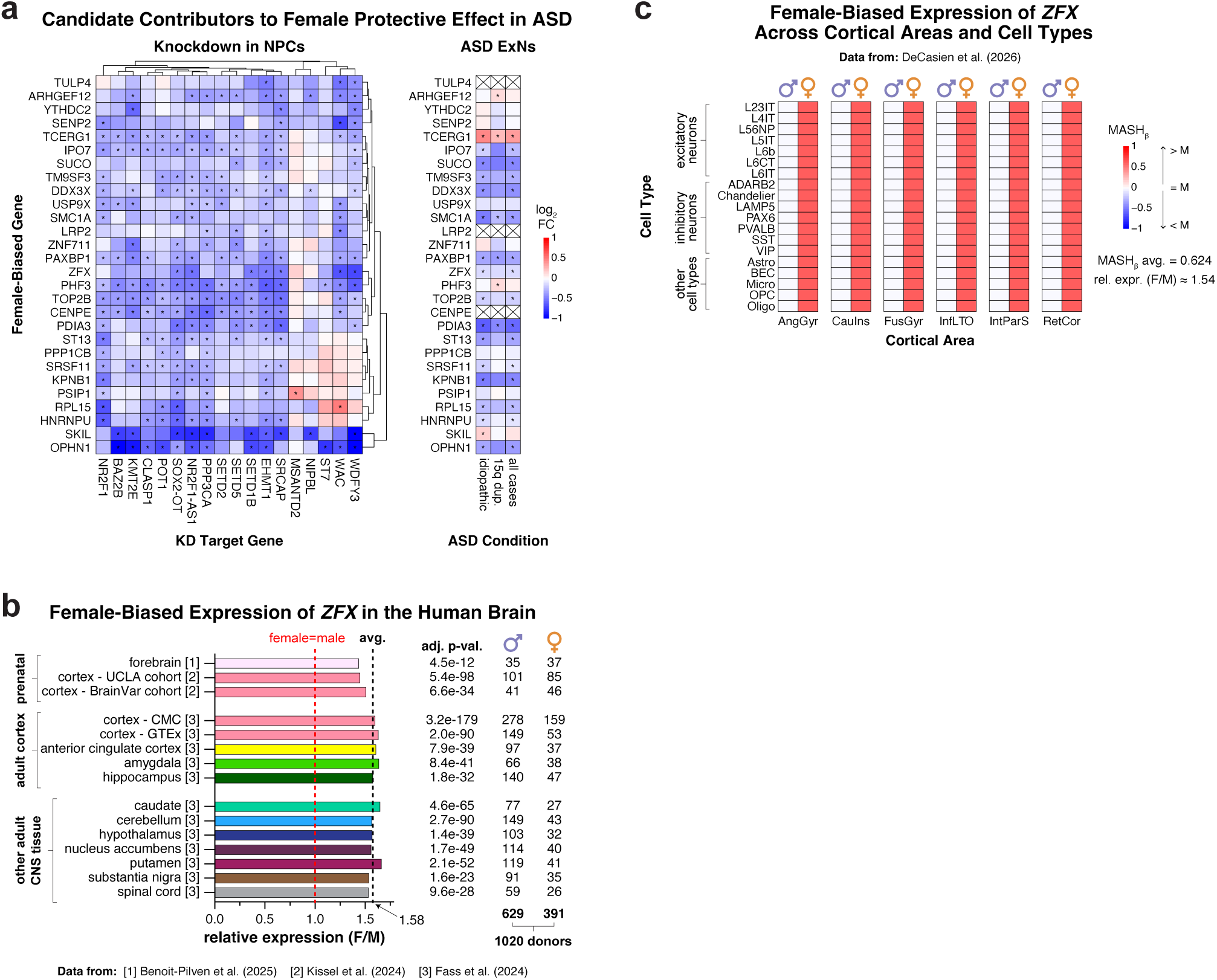
*ZFX* exhibits female-biased expression in the brain and is recurrently downregulated upon ASD gene KD. **a**, Heatmaps showing the relative expression of candidate contributors to the FPE in ASD. Left: expression in NPCs across KD conditions relative to NTCs, with statistically significant DEGs (indicated with *) identified using DESeq2 (Wald test) with Benjamini-Hochberg correction for multiple hypothesis testing. Right: expression in ExNs from males with ASD relative to male idiopathic controls, re-analyzed from Dias et al. (2024)^47^, with statistically significant DEGs (indicated with *) identified using MAST (likelihood ratio test) with Bonferroni correction for multiple hypothesis testing. Lack of detection is indicated with X. **b,** Bar plot depicting the significantly female-biased expression of *ZFX* across several tissues of the human central nervous system (CNS), including prenatal and adult cortex. Distinct regions indicated by different colors. The depicted expression levels and statistical results were taken directly from the following references: [1] Benoit-Pilven et al. (2025)^67^, [2] Kissel et al. (2024)^68^, and [3] Fass et al. (2024)^69^. For details regarding the statistical analyses, see the corresponding reference. **c,** Heatmap depicting expression of *ZFX* across cortical areas and cell types, relative to expression in males. Cell types and areas are abbreviated as in the source, DeCasien et al. (2026)^62^.

## REFERENCES

1. Satterstrom, F. K., et al. Large-Scale Exome Sequencing Study Implicates Both Developmental and Functional Changes in the Neurobiology of Autism Article Large-Scale Exome Sequencing Study Implicates Both Developmental and Functional Changes in the Neurobiology of Autism. Cell 1–17 (2020) doi:10.1016/j.cell.2019.12.036.

2. Trost, B., et al. Genomic architecture of autism from comprehensive whole-genome sequence annotation. Cell 185, 4409–4427.e18 (2022).

3. Ruzzo, E. K., et al. Inherited and De Novo Genetic Risk for Autism Impacts Shared Networks. Cell 178, 850–866.e26 (2019).

4. Liao, C., et al. Convergent coexpression of autism-associated genes suggests some novel risk genes may not be detectable in large-scale genetic studies. Cell Genomics 3, 100277 (2023).

5. Maenner, M. J., et al. Prevalence and Characteristics of Autism Spectrum Disorder Among Children Aged 8 Years - Autism and Developmental Disabilities Monitoring Network, 11 Sites, United States, 2020. MMWR. Surveill. Summ. 72, 1–14 (2023).

6. Jacquemont, S., et al. A higher mutational burden in females supports a ‘female protective model’ in neurodevelopmental disorders. Am. J. Hum. Genet. 94, 415–25 (2014).

7. Antaki, D., et al. A phenotypic spectrum of autism is attributable to the combined effects of rare variants, polygenic risk and sex. Nat. Genet. 54, 1284–1292 (2022).

8. Werling, D. M. The role of sex-differential biology in risk for autism spectrum disorder. Biol. Sex Differ. 7, 58 (2016).

9. Dougherty, J. D., et al. Can the ‘female protective effect’ liability threshold model explain sex differences in autism spectrum disorder? Neuron 110, 3243–3262 (2022).

10. Schneider-Gädicke, A., Beer-Romero, P., Brown, L. G., Nussbaum, R. & Page, D. C. ZFX has a gene structure similar to ZFY, the putative human sex determinant, and escapes X inactivation. Cell 57, 1247–58 (1989).

11. Iakoucheva, L. M., Muotri, A. R. & Sebat, J. Getting to the Cores of Autism. Cell 178, 1287–1298 (2019).

12. Mendes, M., et al. Chromosome X-wide common variant association study in autism spectrum disorder. Am. J. Hum. Genet. 0, (2024).

13. Talukdar, M. & Page, D. C. The inactive X chromosome as a female protector in autism and beyond. Nat. Genet. 58, 687–694 (2026).

14. Abrahams, B. S., et al. SFARI Gene 2.0: a community-driven knowledgebase for the autism spectrum disorders (ASDs). Mol. Autism 4, 36 (2013).

15. Ang, C. E., et al. The novel lncRNA lnc-NR2F1 is pro-neurogenic and mutated in human neurodevelopmental disorders. Elife 8, (2019).

16. Andersen, R. E., et al. Chromosomal structural rearrangements implicate long non-coding RNAs in rare germline disorders. Hum. Genet. 143, 921–938 (2024).

17. Konermann, S., et al. Transcriptome Engineering with RNA-Targeting Type VI-D CRISPR Effectors. Cell 173, 665–676.e14 (2018).

18. Haddad Derafshi, B., et al. The autism risk factor CHD8 is a chromatin activator in human neurons and functionally dependent on the ERK-MAPK pathway effector ELK1. Sci. Rep. 12, 22425 (2022).

19. Hart, S. K., et al. Precise RNA targeting with CRISPR–Cas13d. Nat. Biotechnol. 44, 64–69 (2026).

20. Deneault, E., et al. Complete Disruption of Autism-Susceptibility Genes by Gene Editing Predominantly Reduces Functional Connectivity of Isogenic Human Neurons. Stem Cell Reports 11, 1211–1225 (2018).

21. Fernandez Garcia, M., et al. Transcriptomic and phenotypic convergence of neurodevelopmental disorder risk genes in vitro and in vivo. Nat. Neurosci. 29, 1079–1094 (2026).

22. Gordon, A., et al. Developmental convergence and divergence in human stem cell models of autism. Nature 651, 707–719 (2026).

23. Weerts, M. J. A., et al. Delineating the molecular and phenotypic spectrum of the SETD1B-related syndrome. Genetics in Medicine 23, 2122–2137 (2021).

24. Parra, A., et al. Clinical Heterogeneity and Different Phenotypes in Patients with SETD2 Variants: 18 New Patients and Review of the Literature. Genes (Basel). 14, 1179 (2023).

25. Crippa, M., et al. SETD5 Gene Haploinsufficiency in Three Patients With Suspected KBG Syndrome. Front. Neurol. 11, 631 (2020).

26. O’Donnell-Luria, A. H., et al. Heterozygous Variants in KMT2E Cause a Spectrum of Neurodevelopmental Disorders and Epilepsy. Am. J. Hum. Genet. 104, 1210–1222 (2019).

27. Quental, R., et al. Congenital heart defects associated with pathogenic variants in WAC gene: Expanding the phenotypic and genotypic spectrum of DeSanto-Shinawi syndrome. Am. J. Med. Genet. A 188, 1311–1316 (2022).

28. Connacher, R., et al. Autism NPCs from both idiopathic and CNV 16p11.2 deletion patients exhibit dysregulation of proliferation and mitogenic responses. Stem Cell Reports 17, 1380–1394 (2022).

29. Fombonne, E., Rogé, B., Claverie, J., Courty, S. & Frémolle, J. Microcephaly and macrocephaly in autism. J. Autism Dev. Disord. 29, 113–9 (1999).

30. Nordahl, C. W., et al. Brain enlargement is associated with regression in preschool-age boys with autism spectrum disorders. Proc. Natl. Acad. Sci. U. S. A. 108, 20195–200 (2011).

31. Alsafh, R., et al. Multiplex Consanguineous Family Highlights CLASP1 as a Candidate Gene for Lissencephaly. Neurol. Genet. 10, e200172 (2024).

32. Rots, D., et al. Comprehensive EHMT1 variants analysis broadens genotype-phenotype associations and molecular mechanisms in Kleefstra syndrome. The American Journal of Human Genetics 111, 1605–1625 (2024).

33. Sewani, S., et al. Neurodevelopmental and other phenotypes recurrently associated with heterozygous BAZ2B loss-of-function variants. Am. J. Med. Genet. A 194, e63445 (2024).

34. Paulsen, B., et al. Autism genes converge on asynchronous development of shared neuron classes. Nature 602, 268–273 (2022).

35. Jourdon, A., et al. Modeling idiopathic autism in forebrain organoids reveals an imbalance of excitatory cortical neuron subtypes during early neurogenesis. Nat. Neurosci. 26, 1505–1515 (2023).

36. Voineagu, I., et al. Transcriptomic analysis of autistic brain reveals convergent molecular pathology. Nature 474, 380–4 (2011).

37. Gandal, M. J., et al. Broad transcriptomic dysregulation occurs across the cerebral cortex in ASD. Nature 611, 532–539 (2022).

38. Wamsley, B., et al. Molecular cascades and cell type-specific signatures in ASD revealed by single-cell genomics. Science 384, eadh2602 (2024).

39. Li, C., et al. Single-cell brain organoid screening identifies developmental defects in autism. Nature 621, 373–380 (2023).

40. Schoenherr, C. J. & Anderson, D. J. The neuron-restrictive silencer factor (NRSF): a coordinate repressor of multiple neuron-specific genes. Science 267, 1360–3 (1995).

41. Chong, J. A., et al. REST: a mammalian silencer protein that restricts sodium channel gene expression to neurons. Cell 80, 949–57 (1995).

42. Katayama, Y., et al. CHD8 haploinsufficiency results in autistic-like phenotypes in mice. Nature 537, 675–679 (2016).

43. Abuhatzira, L., Makedonski, K., Kaufman, Y., Razin, A. & Shemer, R. MeCP2 deficiency in the brain decreases BDNF levels by REST/CoREST-mediated repression and increases TRKB production. Epigenetics 2, 214–22 (2007).

44. Tang, X., et al. KCC2 rescues functional deficits in human neurons derived from patients with Rett syndrome. Proc. Natl. Acad. Sci. U. S. A. 113, 751–6 (2016).

45. Sugathan, A., et al. CHD8 regulates neurodevelopmental pathways associated with autism spectrum disorder in neural progenitors. Proc. Natl. Acad. Sci. U. S. A. 111, E4468–77 (2014).

46. Cotney, J., et al. The autism-associated chromatin modifier CHD8 regulates other autism risk genes during human neurodevelopment. Nat. Commun. 6, 6404 (2015).

47. Dias, C., et al. Cell-type-specific effects of autism-associated 15q duplication syndrome in the human brain. Am. J. Hum. Genet. 111, 1544–1558 (2024).

48. San Roman, A. K., et al. The human Y and inactive X chromosomes similarly modulate autosomal gene expression. Cell genomics 4, 100462 (2024).

49. Rhie, S. K., et al. ZFX acts as a transcriptional activator in multiple types of human tumors by binding downstream from transcription start sites at the majority of CpG island promoters. Genome Res. 28, 310–320 (2018).

50. Werling, D. M., Parikshak, N. N. & Geschwind, D. H. Gene expression in human brain implicates sexually dimorphic pathways in autism spectrum disorders. Nat. Commun. 7, 10717 (2016).

51. Velmeshev, D., et al. Single-cell analysis of prenatal and postnatal human cortical development. Science (1979). 382, (2023).

52. Tukiainen, T., et al. Landscape of X chromosome inactivation across human tissues. Nature 550, 244–248 (2017).

53. Shepherdson, J. L., et al. Variants in ZFX are associated with an X-linked neurodevelopmental disorder with recurrent facial gestalt. Am. J. Hum. Genet. 111, 487–508 (2024).

54. Chahrour, M. H., et al. Whole-exome sequencing and homozygosity analysis implicate depolarization-regulated neuronal genes in autism. PLoS Genet. 8, e1002635 (2012).

55. Zhou, X., et al. Integrating de novo and inherited variants in 42,607 autism cases identifies mutations in new moderate-risk genes. Nat. Genet. 54, 1305–1319 (2022).

56. Czerminski, J. T., King, O. D. & Lawrence, J. B. Large-scale organoid study suggests effects of trisomy 21 on early fetal neurodevelopment are more subtle than variability between isogenic lines and experiments. Front. Neurosci. 16, 972201 (2022).

57. Zyla, J., et al. Evaluation of zero counts to better understand the discrepancies between bulk and single-cell RNA-Seq platforms. Comput. Struct. Biotechnol. J. 21, 4663–4674 (2023).

58. Chatterjee, S. et al. Enhancer Variants Synergistically Drive Dysfunction of a Gene Regulatory Network In Hirschsprung Disease. Cell 167, 355–368.e10 (2016).

59. Yim, K. M., et al. Cell-type-specific dysregulation of gene expression due to Chd8 haploinsufficiency during mouse cortical development. Cell genomics 5, 100986 (2025).

60. Durak, O., et al. Chd8 mediates cortical neurogenesis via transcriptional regulation of cell cycle and Wnt signaling. Nat. Neurosci. 19, 1477–1488 (2016).

61. Willsey, H. R., et al. Parallel in vivo analysis of large-effect autism genes implicates cortical neurogenesis and estrogen in risk and resilience. Neuron 109, 788–804.e8 (2021).

62. DeCasien, A. R., et al. Sex effects on gene expression across the human cerebral cortex at cell type resolution. Science 392, eaea9063 (2026).

63. Ferrer, J. & Dimitrova, N. Transcription regulation by long non-coding RNAs: mechanisms and disease relevance. Nat. Rev. Mol. Cell Biol. (2024) doi:10.1038/s41580-023-00694-9.

64. Wong, C. C. Y., et al. Genome-wide DNA methylation profiling identifies convergent molecular signatures associated with idiopathic and syndromic autism in post-mortem human brain tissue. Hum. Mol. Genet. 28, 2201–2211 (2019).

65. Weiner, D. J., et al. Statistical and functional convergence of common and rare genetic influences on autism at chromosome 16p. Nat. Genet. 54, 1630–1639 (2022).

66. Rockowitz, S. & Zheng, D. Significant expansion of the REST/NRSF cistrome in human versus mouse embryonic stem cells: potential implications for neural development. Nucleic Acids Res. 43, 5730–43 (2015).

67. Benoit-Pilven, C., et al. Early establishment and life course stability of sex biases in the human brain transcriptome. Cell genomics 5, 100890 (2025).

68. Kissel, L. T., et al. Sex-Differential Gene Expression in Developing Human Cortex and Its Intersection With Autism Risk Pathways. Biological psychiatry global open science 4, 100321 (2024).

69. Fass, S. B., et al. Relationship between sex biases in gene expression and sex biases in autism and Alzheimer’s disease. Biol. Sex Differ. 15, 47 (2024).

70. Wei, J., et al. Deep learning and CRISPR-Cas13d ortholog discovery for optimized RNA targeting. Cell Syst. 14, 1087–1102.e13 (2023).

71. Camacho, C. et al. BLAST+: architecture and applications. BMC Bioinformatics 10, 421 (2009).

72. Ventura, A., et al. Cre-lox-regulated conditional RNA interference from transgenes. Proc. Natl. Acad. Sci. U. S. A. 101, 10380–5 (2004).

73. Yusa, K., Zhou, L., Li, M. A., Bradley, A. & Craig, N. L. A hyperactive piggyBac transposase for mammalian applications. Proc. Natl. Acad. Sci. U. S. A. 108, 1531–6 (2011).

74. Kumamoto, T., et al. Direct Readout of Neural Stem Cell Transgenesis with an Integration-Coupled Gene Expression Switch. Neuron 107, 617–630.e6 (2020).

75. Schnütgen, F., et al. A directional strategy for monitoring Cre-mediated recombination at the cellular level in the mouse. Nat. Biotechnol. 21, 562–5 (2003).

76. Kaczmarczyk, S. J. & Green, J. E. A single vector containing modified cre recombinase and LOX recombination sequences for inducible tissue-specific amplification of gene expression. Nucleic Acids Res. 29, E56–6 (2001).

77. Qian, K., et al. A simple and efficient system for regulating gene expression in human pluripotent stem cells and derivatives. Stem Cells 32, 1230–8 (2014).

78. Schlaeger, T. M., et al. A comparison of non-integrating reprogramming methods. Nat. Biotechnol. 33, 58–63 (2015).

79. de Leeuw, V. C., et al. An efficient neuron-astrocyte differentiation protocol from human embryonic stem cell-derived neural progenitors to assess chemical-induced developmental neurotoxicity. Reprod. Toxicol. 98, 107–116 (2020).

80. Giandomenico, S. L., Sutcliffe, M. & Lancaster, M. A. Generation and long-term culture of advanced cerebral organoids for studying later stages of neural development. Nat. Protoc. 16, 579–602 (2021).

81. Han, X., et al. Millisecond-timescale optical control of neural dynamics in the nonhuman primate brain. Neuron 62, 191–8 (2009).

82. Lohman, B. K., Weber, J. N. & Bolnick, D. I. Evaluation of TagSeq, a reliable low-cost alternative for RNAseq. Mol. Ecol. Resour. 16, 1315–1321 (2016).

83. Rabe, B. A. Investigation of the Multiple Roles of Notch Signaling in Embryonic Retinal Development Using Novel High-Throughput Techniques. (Harvard University Graduate School of Arts and Sciences, 2020).

84. Ye, J., et al. Primer-BLAST: a tool to design target-specific primers for polymerase chain reaction. BMC Bioinformatics 13, 134 (2012).

85. Kochinke, K., et al. Systematic Phenomics Analysis Deconvolutes Genes Mutated in Intellectual Disability into Biologically Coherent Modules. Am. J. Hum. Genet. 98, 149–64 (2016).

86. Karczewski, K. J., et al. The mutational constraint spectrum quantified from variation in 141,456 humans. Nature 581, 434–443 (2020).

87. Wang, T., et al. Integrated gene analyses of de novo variants from 46,612 trios with autism and developmental disorders. Proceedings of the National Academy of Sciences 119, e2203491119 (2022).

88. Doan, R. N., et al. Recessive gene disruptions in autism spectrum disorder. Nat. Genet. 51, 1092–1098 (2019).

89. Rodin, R. E., et al. The landscape of somatic mutation in cerebral cortex of autistic and neurotypical individuals revealed by ultra-deep whole-genome sequencing. Nat. Neurosci. 24, 176–185 (2021).

90. Wilfert, A. B., et al. Recent ultra-rare inherited variants implicate new autism candidate risk genes. Nat. Genet. 53, 1125–1134 (2021).

91. Alcantara, D., et al. Mutations of AKT3 are associated with a wide spectrum of developmental disorders including extreme megalencephaly. Brain 140, 2610–2622 (2017).

92. Marcelis, A. & Van Reet, E. A Boy With KIF11-Associated Disorder Along With ADHD and ASD: Collaboration Between Paediatrics and Child Psychiatry. Case Rep. Psychiatry 2024, 5535830 (2024).

93. Ruiz-Reig, N., Hakanen, J. & Tissir, F. Connecting neurodevelopment to neurodegeneration: a spotlight on the role of kinesin superfamily protein 2A (KIF2A). Neural Regen. Res. 19, 375–379 (2024).

94. McGhee, C. A., et al. Genotype-phenotype correlations with autism spectrum disorder-related traits in noonan syndrome and noonan syndrome with multiple lentigines: a cross-sectional study. Mol. Autism 16, 51 (2025).

95. Roopra, A. MAGIC: A tool for predicting transcription factors and cofactors driving gene sets using ENCODE data. PLoS Comput. Biol. 16, e1007800 (2020).

96. Lachmann, A., Giorgi, F. M., Lopez, G. & Califano, A. ARACNe-AP: gene network reverse engineering through adaptive partitioning inference of mutual information. Bioinformatics 32, 2233–2235 (2016).

97. Lambert, S. A., et al. The Human Transcription Factors. Cell 172, 650–665 (2018).

98. Boukas, L., et al. Coexpression patterns define epigenetic regulators associated with neurological dysfunction. Genome Res. 29, 532–542 (2019).

99. Alvarez, M. J., et al. Functional characterization of somatic mutations in cancer using network-based inference of protein activity. Nat. Genet. 48, 838–47 (2016).

100. Page, L., Brin, S., Motwani, R. & Winograd, T. The PageRank Citation Ranking: Bringing Order to the Web. The Web Conference https://www.semanticscholar.org/paper/The-PageRank-Citation-Ranking-%3A-Bringing-Order-to-Page-Brin/eb82d3035849cd23578096462ba419b53198a556 (1999).

101. Subramanian, A., et al. Gene set enrichment analysis: A knowledge-based approach for interpreting genome-wide expression profiles. Proceedings of the National Academy of Sciences 102, 15545–15550 (2005).

102. Liberzon, A., et al. The Molecular Signatures Database (MSigDB) hallmark gene set collection. Cell Syst. 1, 417–425 (2015).

103. Szklarczyk, D., et al. The STRING database in 2023: protein-protein association networks and functional enrichment analyses for any sequenced genome of interest. Nucleic Acids Res. 51, D638–D646 (2023).

104. Oliva, M., et al. The impact of sex on gene expression across human tissues. Science 369, (2020).

105. Naqvi, S., et al. Conservation, acquisition, and functional impact of sex-biased gene expression in mammals. Science 365, (2019).

106. Finak, G., et al. MAST: a flexible statistical framework for assessing transcriptional changes and characterizing heterogeneity in single-cell RNA sequencing data. Genome Biol. 16, 278 (2015).

