## Supplementary material for "Transcriptomic Convergence and the Female Protective Effect in Autism": Guide to Supplementary Information

**Supplementary Figure:** Representative flow cytometry gating strategy for analysis of CytoTrack proliferation assays in NPCs, corresponding to **Fig. 1f** and **Extended Data Fig. 6a**.

**Supplementary Table 1:** Guide RNA (gRNA) sequences used with CasRx.

**Supplementary Table 2:** Data associated with **Extended Data Fig. 1a**.

**Supplementary Table 3:** Data associated with **Fig. 1c**.

**Supplementary Table 4:** Normalized expression levels of fluorophores in comparison to ASD gene targets, associated with **Extended Data Fig. 4b**.

**Supplementary Table 5:** Data associated with **Extended Data Fig. 4b**.

**Supplementary Table 6:** Adj. p values for pairwise cross-enrichment of downregulated DEGs, associated with **Extended Data Fig. 5**.

**Supplementary Table 7:** Median LOEUF scores used for assessing constraint. Highly constrained genes were defined as having a median LOEUF score of less than 0.35.

**Supplementary Table 8:** Genes that meet the criteria used for identifying candidate ASD risk genes: 1) statistically significant downregulation by at least three of the ASD KDs, with at least 80% concordance in the direction of significant differential expression across all KDs (consistently downregulated DEGs); and 2) intolerance to loss of function variants (highly constrained genes).

**Supplementary Table 9:** Candidate ASD risk genes from **Supplementary Table 8** that also show some association with ASD in one or more major sequencing studies[2,55,87–90], referenced by PMID.

**Supplementary Table 10:** Full statistical results associated with CytoTrack NPC proliferation assays depicted in **Fig. 1f** and **Extended Data Fig. 6a**.

**Supplementary Table 11:** Top significant MAGIC regulators identified in different cell populations. For each population, MAGIC regulators were scored by the number of ASD gene KDs that implicated the regulator. The 25 top scoring regulators are included here, along with any additional regulators with a tied score. Associated with **Extended Data Fig. 8a**.

**Supplementary Table 12:** Top drivers of convergence in ASD, as identified through GRN analysis. Associated with **Fig. 4e**.

**Supplementary Table 13:** Sequences of shRNAs.

**Supplementary Table 14:** Oligo sequences used to generate libraries for bulk RNA-sequencing.

**Supplementary Table 15:** Primers for ddPCR/qPCR.

**Supplementary Tables 16-35:** Differential expression results following CasRx-based KD in NPCs.

**Supplementary Tables 36-41:** Differential expression results following shRNA-based KD in NPCs.

**Supplementary Tables 42-61**: Differential expression results from re-analysis of CRISPR-Cas9 KO of ASD genes in iPSCs and ExNs, from Deneault et al. (2018)[20].

**Supplementary Tables 62-64:** Differential expression results following CasRx-based KD in mosaic organoids, as analyzed by scRNA-seq.

**Supplementary Table 65:** Differential expression results from re-analysis of snRNA-seq data from ASD cases vs controls, from Dias et al. (2024)[47].
