## Supplementary Figure for "Transcriptomic Convergence and the Female Protective Effect in Autism"

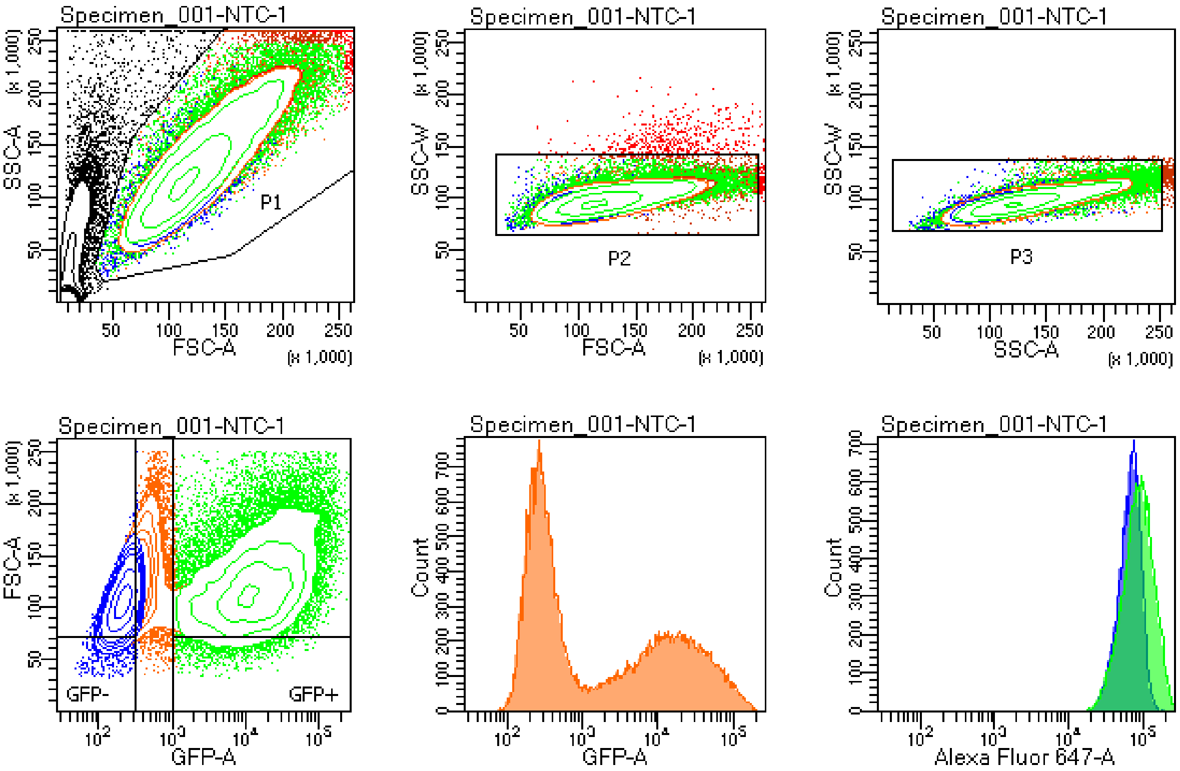


Tube: NTC-1

| Population | #Events | %Parent | %Total |
|---|---|---|---|
| All Events | 58,990 | #### | 100.0 |
| P1 | 44,418 | 75.3 | 75.3 |
| P2 | 43,045 | 96.9 | 73.0 |
| P3 | 42,168 | 98.0 | 71.5 |
| GFP- | 15,431 | 36.6 | 26.2 |
| GFP+ | 19,911 | 47.2 | 33.8 |

| Population | #Events | %Parent | Alexa Flu... Geo Mean |
|---|---|---|---|
| P1 | 44,418 | 75.3 | 76,721 |
| P2 | 43,045 | 96.9 | 74,700 |
| P3 | 42,168 | 98.0 | 73,352 |
| GFP- | 15,431 | 36.6 | 60,972 |
| GFP+ | 19,911 | 47.2 | 78,927 |

**Supplementary Figure:** Representative flow cytometry gating strategy for analysis of CytoTrack proliferation assays in NPCs, corresponding to **Fig. 1f** and **Extended Data Fig. 6a**.
